# cGAS recruitment to micronuclei is dictated by pre-existing nuclear chromatin status

**DOI:** 10.1101/2022.01.13.476191

**Authors:** Kate M. MacDonald, Shirony Nicholson-Puthenveedu, Maha M. Tageldein, Cheryl Arrowsmith, Shane M. Harding

## Abstract

Micronuclei (MN) are aberrant cytosolic compartments containing broken genomic fragments or whole lagging chromosomes. MN envelopes irreversibly rupture, allowing the viral receptor cGAS to localize to MN and initiate an inflammatory signalling cascade. Here, we demonstrate that MN envelope rupture is not sufficient for cGAS localization. Unlike MN that arise following ionizing radiation (IR), ruptured MN generated from acute transcription stressors DRB or siSRSF1 are refractory to cGAS localization. Recruitment of cGAS to MN is blocked by inhibiting the histone methyltransferase DOT1L prior to IR exposure, demonstrating that cGAS recruitment to MN is dictated by nuclear chromatin organization at the time of DNA damage. Loss of cGAS+ MN, caused either by acute transcription stressors or by preventing DOT1L-deposited histone methylation, corresponded to significantly decreased cGAS-dependent inflammatory signalling. These results implicate nuclear chromatin organization in micronuclear composition and activity, influencing the ability of damage-induced MN to retain cytosolic proteins upon rupture.

## INTRODUCTION

Micronuclei (MN) contain pieces of the nuclear genome, sequestered in the cytosol following mitosis where there is unresolved DNA damage (Terradas et al., 2010; MacDonald, Benguerfi and Harding, 2020). MN arise at mitotic exit, when acentric chromosomal fragments or whole lagging chromosomes fail to be partitioned into newly forming daughter nuclei. Instead, these fragments recruit their own nuclear envelopes, discontinuous with the primary nuclear membrane. MN are active participants in carcinogenic processes including chromothripsis and double-stranded DNA (dsDNA)-driven inflammatory cascades, that are in part dependent on the integrity of the micronuclear envelope (Terradas et al., 2010; Crasta et al., 2012; Zhang et al., 2015; Terradas, Martín and Genescà, 2016; Harding et al., 2017; Mackenzie et al., 2017; Umbreit et al., 2020). Compared to primary nuclei, MN envelopes tend to be structurally defective, predisposed to irreparable breaches, called rupture, in interphase (Hatch et al., 2013; S. Liu et al., 2018; Vietri et al., 2020). Rupture exposes dsDNA to the cytosol, where it recruits at least one sentinel viral pattern recognition receptor (PRR), called cyclic GMP-AMP (cGAMP) synthase (cGAS) (Harding et al., 2017; Mackenzie et al., 2017). Canonically, cGAS recognizes invading dsDNA viruses and produces cGAMP, which activates endoplasmic reticulum-bound stimulator of interferon genes (STING). STING initiates a cascade of anti-viral responses, alerting the immune system to a possible viral infection (Li et al., 2013; Abe and Barber, 2014). The same cGAS-STING activity is induced after DNA damage, eliciting expression of a suite of inflammatory genes and cytokines following recognition of ruptured MN by cGAS (Gao et al., 2013; Li et al., 2013; Abe and Barber, 2014; Harding et al., 2017).

MN rupture dynamics can be affected by the identity of an enclosed whole chromosome (Sepaniac et al., 2021; Mammel et al., 2022). MN DNA content, whether a whole chromosome or an acentric chromosomal fragment, is influenced by the location of the inciting genomic insult (Ly et al., 2017; Leibowitz et al., 2021). To date, work on MN envelope dynamics and cGAS recognition has focused primarily on MN generated from double-stranded break (DSB)-inducing IR exposure (Harding et al., 2017; Mackenzie et al., 2017) or MN containing whole lagging chromosomes after mitotic spindle poison (Crasta et al., 2012; Hatch et al., 2013; Zhang et al., 2015; Umbreit et al., 2020; Vietri, et al., 2020; Flynn, Koch and Mitchison, 2021; Mohr et al., 2021; Mammel et al., 2022) both of which are expected to generate MN with essentially randomized genomic content. Under these conditions, MN rupture appears to be a primary determinant of cGAS access (Harding et al., 2017; Mackenzie et al., 2017; Mohr et al., 2021). MN have also been observed following cellular exposure to more common and targeted forms of DNA damage, including alkylating base-damaging agents (Lutz et al., 2005) and acute transcription stressors (Gan et al., 2011), which deposit lesions that can transition to DSBs. These different classes of genotoxic stressors target different regions of the genome, potentially impacting the location of unresolved DSBs (Volkova et al., 2020). As such, the inciting genotoxic exposure could influence MN composition and structure by dictating MN content. Changes to MN content and composition could affect rupture dynamics and cGAS retention, and the downstream inflammatory cascade.

Here, we ask whether acute cellular exposure to distinct genotoxic agents influences the characteristics of the micronuclei produced. We find that cGAS recognition of MN depends on the DNA-damaging agent to which the cell was initially exposed. Surprisingly, MN envelope rupture was not sufficient for cGAS to localize. Unlike MN formed following IR exposure, alkylating base damage, or treatment with an aneugenic agent, the MN produced following acute transcription stressors were almost entirely cGAS-negative, even when ruptured. Our data suggest that cGAS recognition of MN DNA depends on the chromatin structure of the sequestered fragment, rather than being purely dictated by access through the MN envelope. cGAS binding to IR-induced MN could be manipulated with pharmacological inhibitors of histone modifying enzymes, particularly the histone 3 lysine 79 (H3K79) methyltransferase disruptor of telomere silencing 1-like (DOT1L), or by forcing active transcription at the damage site. Loss of cGAS+ MN, caused either by acute transcription stressors or by preventing nuclear H3K79 dimethylation, corresponded to significantly decreased cGAS-dependent interferon stimulated gene (ISG) expression. Together these data suggest that micronuclear heterogeneity is predetermined by the state of the enclosed chromatin, with important implications for the emergence of genomic instability and tumorigenesis and the response to genotoxic cancer therapies.

## RESULTS

### cGAS recognition of micronuclei depends on the inciting genotoxic stressor

Micronuclei can be visualized and scored as cGAS+ or cGAS-by immunofluorescent microscopy (IF) (**Fig. 1A**) or by flow cytometry (Mohr et al., 2021) (**Fig. S1A**). We acutely exposed MCF10A cells to a panel of distinct genotoxic stressors, removed the treatments, and allowed 72 hours for the cells to divide and produce MN (**Fig. 1B**). Doses and exposure times for each agent were chosen to maximize MN production while minimizing cell death within the time frame of the assay (see *Methods*). At 72 hours post-acute exposure, the percentage of cGAS+ MN were counted by IF, demonstrating a significant association with the inciting genotoxic agent (**Fig. 1C**). This phenomenon was also observed in HeLa and U2OS cells and confirmed with flow cytometry in HeLa (**Fig. S1B**). At the extremes of the % cGAS+ MN distribution were IR, a nonspecific clastogen previously shown to generate cGAS-bound MN (Harding et al., 2017; Mackenzie et al., 2017); methyl methanesulfonate (MMS), a base-alkylating agent; 5,6-dichloro-1-beta-ribo-furanosyl benzimidazole (DRB), an RNA polymerase II inhibitor; and an siRNA-mediated knockdown of serine/arginine-rich splicing factor 1 (SRSF1), a pre-mRNA splicing protein. All of these are expected to generate MN containing acentric chromosomal fragments, as supported by centromere protein A (CENP-A) staining in the MN (**Fig. S1C**). Along with paclitaxel (Taxol), which induces whole chromosome mis-segregation into MN, this reduced panel of genotoxic agents was selected for continuing experiments on MN-cGAS recruitment.

**Figure 1.**
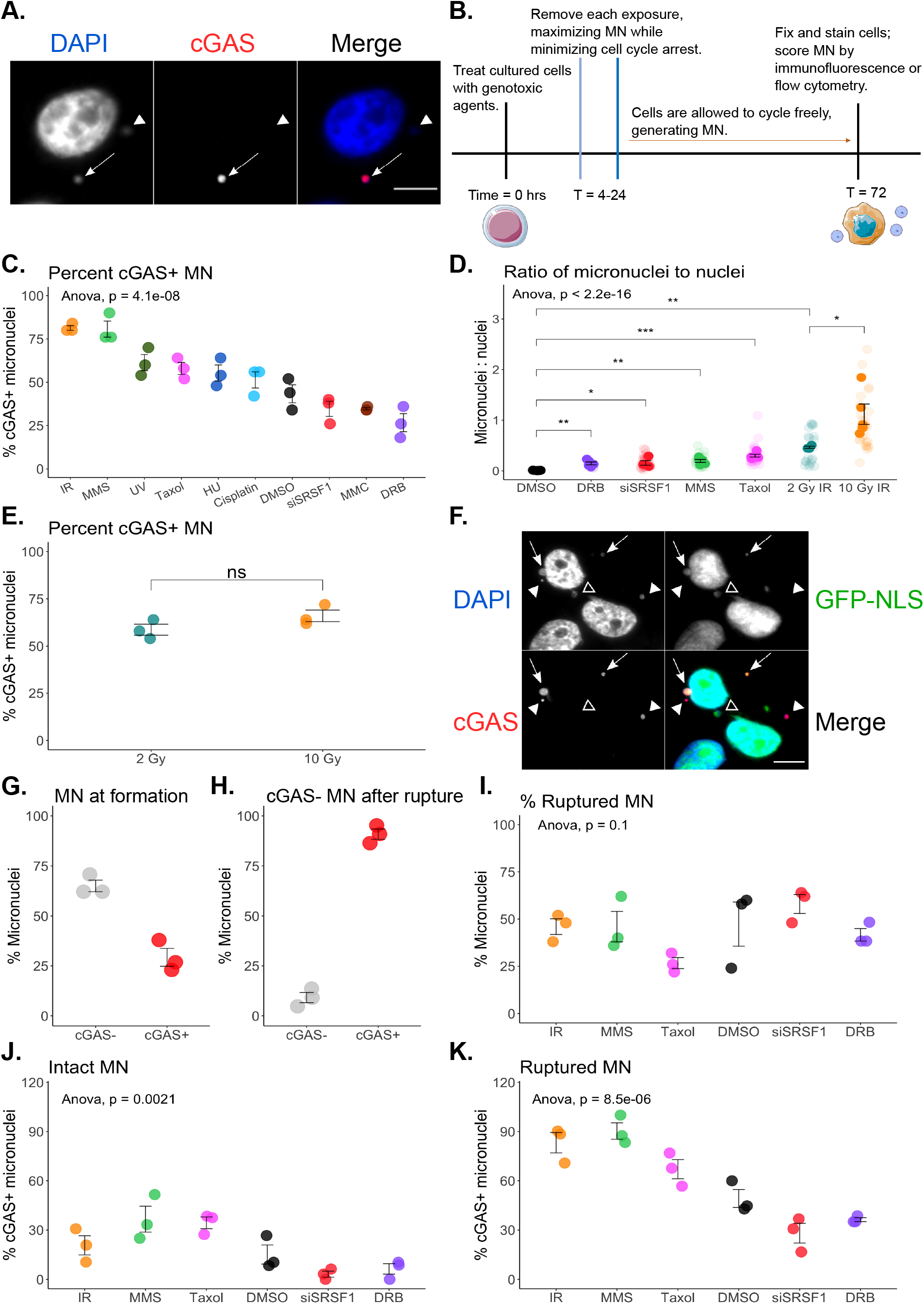
cGAS recognition of micronuclei depends on the inciting genotoxic stressor, and micronuclear envelope rupture is not sufficient for cytosolic cGAS recognition. (**A**) MCF10A cells 72 hours post-IR exposure. Arrow = cGAS+ MN, arrowhead = cGAS-MN. Scale bar = 10 μm. (**B**) Timeline (hours) for immunofluorescence (IF) experiments used to score MN. (**C**) Percent cGAS+ MN observed by IF, 72 hours post-acute exposure. *N* per biological replicate = 50 MN. Statistical comparison by two-way ANOVA. (**D**) Observed ratio of MN to primary nuclei, 72 hours post-acute exposure. *N* per biological replicate = 5-8 fields of view. Statistical comparison by Welch’s t-test. (**E**) Percent cGAS+ MN observed by IF, 72 hours following exposure to 2 or 10 Gy IR. *N* per biological replicate = 50 MN. Statistical comparison by Student’s t-test. (**F**) MCF10A cells expressing a GFP-tagged nuclear localization signal (GFP-NLS), 72 hours post-IR exposure. Arrow = GFP+ cGAS+ MN (intact, cGAS+), arrowhead = GFP-cGAS+ MN (ruptured, cGAS+), empty arrowhead = GFP+ cGAS-MN (intact, cGAS-MN). Scale bar = 10 μm. (**G**) Percent IR-induced MN that are cGAS-vs. cGAS+ at the time of their formation. *N* per biological replicate = 50 MN. (**H**) Within the subset of IR-induced MN that were cGAS-at the time of their formation, the percent MN that acquired cGAS within one hour of rupture. *N* per biological replicate = 50 MN. (**I**) Percent ruptured MN observed by IF, 72 hours post-acute exposure. *N* per biological replicate = 50 MN. Statistical comparison by twoway ANOVA. (**J**) Percent intact, cGAS+ MN observed by IF, 72 hours post-acute exposure. *N* per biological replicate = 50 MN. Statistical comparison by two-way ANOVA. (**K**) Percent ruptured, cGAS+ MN observed by IF, 72 hours post-acute exposure. *N* per biological replicate = 50 MN. Statistical comparison by two-way ANOVA. ns: *p* > 0.05, *: *p*<= 0.05, **: *p*<= 0.01, ***: *p*<= 0.001, ****: *p*<= 0.0001. Data are represented as mean ± SEM. See also Figure S1, Figure S2.

Divergent cGAS recruitment to MN reflects either differences intrinsic to the MN, or variable effects on cellular cGAS protein, following acute genotoxic stressors. Acute treatments did not affect cGAS RNA or protein levels, meaning changes to cGAS expression are not responsible for differential % cGAS+ MN (**Fig. S2A-B**). Each acute exposure generated a different number of MN per primary nucleus (**Fig. 1D**). This outcome might be expected following differences in the absolute number of unresolved DSBs across treatments. In all cases, we observed a high correlation between γH2AX+ (a marker of DSBs) and cGAS+ MN (**Fig. S2C**, *R*^2^ = 0.88) (H. Liu et al., 2018), though we did not detect a direct interaction between cGAS and γH2AX by co-immunoprecipitation (**Fig. S2D**). Despite this correlation with γH2AX, we found no relationship between the proportion of cGAS+ MN and the total number of MN produced after each acute treatment (**Fig. S2E**, *R*^2^ = 0.41). Explicitly manipulating the number of DSBs and MN with two different doses of ionizing radiation did not affect % cGAS+ MN (2 Gy and 10 Gy; **Fig. 1E**). Thus, despite favouring γH2AX-staining regions, cGAS recruitment to MN appears independent of the total damage deposited by each agent. Finally, it has been previously shown that cGAS localization depends in part on cell cycle phase (Harding et al., 2017; Li et al., 2021). Mitotic cells can inhibit cGAS via phosphorylation and nuclear chromatin tethering (Li et al., 2021). Therefore, cells that spend more time in interphase might be afforded greater opportunity for cGAS recognition of MN. We asked whether alterations in cell cycle progression induced by each acute genotoxic exposure could explain their differential % cGAS+ MN, using IR (high % cGAS+ MN) and siSRSF1 (low % cGAS+ MN) as test cases, in mCherry-H2B MCF10A cells followed by live-cell imaging. After acute IR or siSRSF1, the overall population growth curves were different (**Fig. S2F**) but the average time that a micronucleated cell spent in interphase was unaffected (**Fig. S2G**). Together these findings suggest that cGAS localization to MN across acute genotoxic exposures is determined by intrinsic features of MN, rather than changes to cGAS protein expression or cGAS tethering to mitotic chromatin.

### Micronuclear envelope rupture is not sufficient for cytosolic cGAS recognition

cGAS recognition of MN has been attributed to micronuclear envelope rupture (Mackenzie et al., 2017; Mohr et al., 2021). To assess whether the observed difference in % cGAS+ MN across genotoxic exposures could be explained by differences in MN envelope stability, we used MCF10A cells expressing a GFP-tagged nuclear localization signal (NLS). The GFP-NLS tag is only found inside intact MN, and diffusion of GFP upon MN rupture is a reliable indicator of MN envelope integrity (Hatch et al., 2013; Mackenzie et al., 2017; Mohr et al., 2021). To avoid any confounding effects on cGAS MN localization coming from nuclear cGAS, we relied on MCF10A cells for cytosolic MN rupture experiments, where cGAS is primarily a cytosolic protein at baseline (**Fig. S2H**). At 72 hours post-acute genotoxic exposure, MN can be scored as cGAS+ intact (cGAS+, GFP-NLS+), cGAS+ ruptured (cGAS+, GFP-), cGAS-intact, or cGAS-ruptured by immunofluorescence (**Fig. 1F**). Consistent with published reports, we found using live-cell imaging that following 10 Gy IR exposure, ~75% of MN initially form as cGAS-(**Fig. 1G**) and that when cGAS-MN rupture in the cytosol, ~90% of those MN acquire cGAS within one hour (**Fig. 1H**) (Mackenzie et al., 2017; Mohr et al., 2021).

We expected that the MN generated following each acute genotoxic stressor in our panel would demonstrate variance in their envelope integrity, where high incidence of ruptured MN would correlate with high cGAS+ MN. Instead, we observed no significant difference in % ruptured MN at 72 hours post-exposure (**Fig. 1I**). Percent cGAS recognition of both intact and ruptured MN was dependent on the genotoxic agent to which the cell had been exposed (**Fig. 1J-K**). In spontaneously occurring MN (DMSO control) 49% of ruptured MN recruit cGAS, while those produced by IR or MMS exposure recruit at cGAS 82% and 90% incidence, respectively (**Fig. 1K**). After siSRSF1 or DRB, both agents that interfere with transcription, ruptured MN were recognized by cGAS in just 28% and 36% of cases, respectively (**Fig. 1K**). The endoplasmic reticulum (ER) is recruited to MN upon rupture, and brings ER-associated factors including the nuclease TREX1, which has been reported as a negative regulator of cGAS (Hatch et al., 2013; Vietri, et al., 2020; Mohr et al., 2021). We observed no correlating difference in % MN associated with CHMP7 (a marker of ongoing ER-MN invasion (Vietri et al., 2020)) across exposures, suggesting that ER-associated factors are not responsible for the variance in cGAS recruitment to ruptured MN in this panel (**Fig. S2I**). Together, these data show that rupture of MN alone is not sufficient for cGAS retention. Our results suggest the existence of other MN-intrinsic elements, varying across genotoxic exposures, that affect cGAS recruitment upon cytosolic MN envelope rupture.

### cGAS does not bind MN originating from ongoing, active transcription

DRB or siSRSF1-induced MN rupture at the same frequency as the MN produced by IR (**Fig. 1I**), but they are refractory to cGAS (**Fig. 1K**). Both DRB and siSRSF1 are acute transcription stressors, and we asked whether cellular exposure to DRB or siSRSF1 was enriching MN content for transcription-associated units. We again evaluated cGAS+ and cGAS-MN 72 hours postexposure, co-staining cells with 5-ethynyl uridine (EU), a fluorescent marker of ongoing transcription, one hour prior to collecting cells for IF microscopy. We found cGAS and EU to be mutually exclusive in MN across all agents in both MCF10A and HeLa cells, suggesting cGAS is avoiding actively transcribing regions in MN (**Fig. 2A-C, Fig. S3A**). MN lose their capacity to transcribe when their envelope ruptures (Hatch et al., 2013) and since ruptured MN recruit cGAS (**Fig. 1J-K**), we first considered that the dichotomy between cGAS+ and EU+ MN was a function of envelope integrity. However, MN envelope integrity does not account for differences in cGAS recruitment across agents, including DRB and siSRSF1 (**Fig. 1I-K**), while EU+ status strongly anticorrelates (**Fig. 2C**). Furthermore, on the rare occasion when a micronucleus was both EU+ and cGAS+, the two markers did not spatially overlap, demonstrating a discrete pattern even when both cGAS and transcription machinery had equal access (**Fig. 2A, bottom row**). These findings suggest that transcription-associated chromatin features present at the time of damage induction may dissuade cGAS localization to otherwise accessible MN.

**Figure 2.**
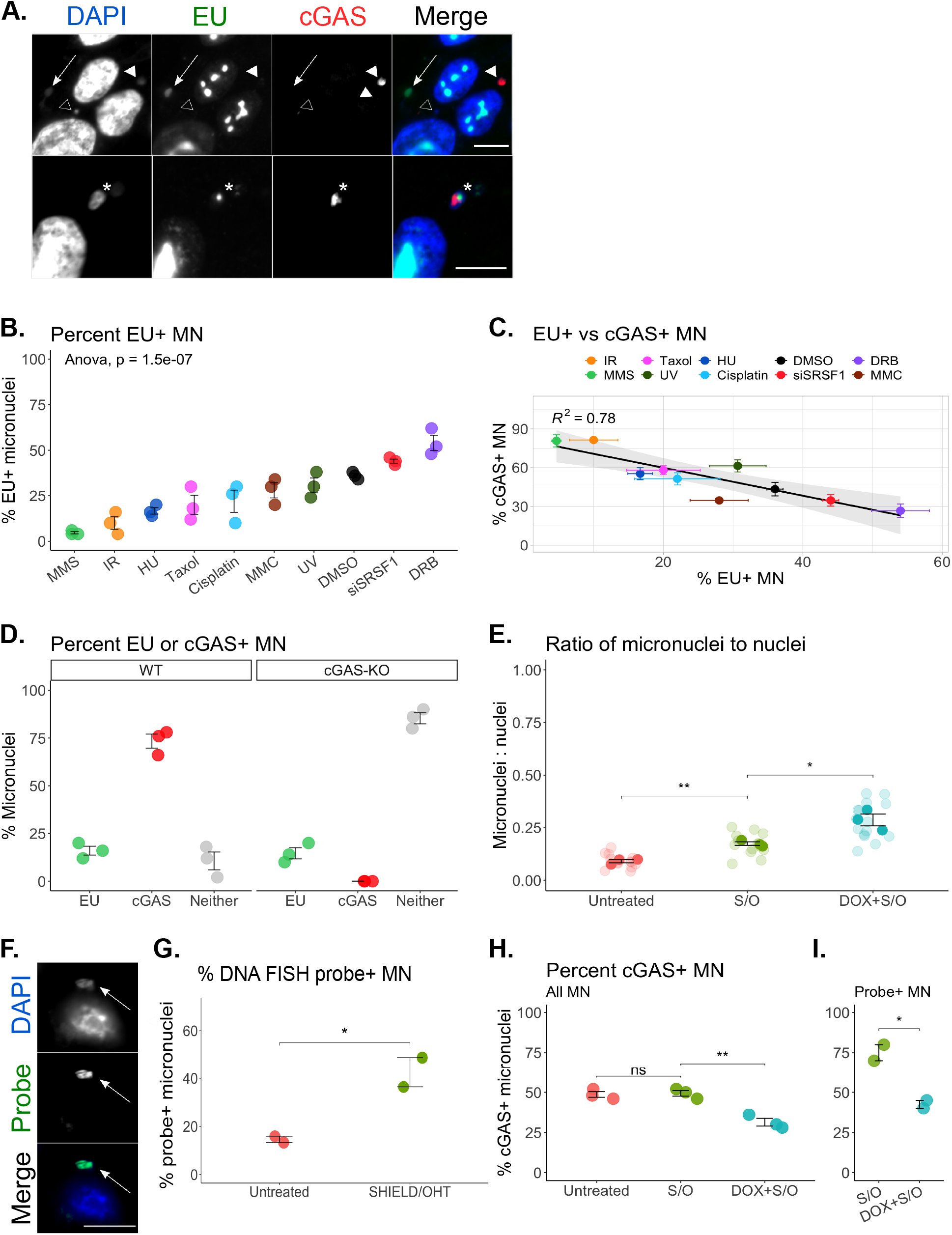
cGAS does not bind MN originating from ongoing, active transcription. (**A**) MCF10A cells 72 hours post-IR exposure. Arrow = EU+ MN, arrowhead = cGAS+ MN, empty arrowhead = EU-cGAS-MN, asterisk = EU+ cGAS+ MN. Scale bar = 20 μm. (**B**) Percent EU+ MN observed by IF, 72 hours post-acute exposure. *N* per biological replicate = 50 MN. Statistical comparison by two-way ANOVA. (**C**) Linear regression comparing the % EU+ to the % cGAS+ MN observed by IF, 72 hours post-acute exposure. Each data point represents the mean of three biological replicates, where 50 MN were counted per replicate. (**D**) Percent EU+, cGAS+, or neither+ MN observed by IF, 72 hours post-acute 10 Gy IR, in wild-type (WT) vs. cGAS-knockout MCF10A cells. *N* per biological replicate = 50 MN. (**E**) Observed ratio of MN to primary nuclei, 72 hours post-acute SHIELD/OHT. *N* per biological replicate = 5 fields of view. Statistical comparison by Student’s t-test. (**F**) DNA FISH with SHIELD/OHT-treated 256 U2OS cells to verify that SHIELD/OHT-targeted region of the genome is sequestered in the resulting MN. Arrow = FISH probe+ MN. Scale bar = 10 μm. (**G**) Percent FISH probe+ MN observed by IF, 72 hours post-acute exposure to SHIELD/OHT. *N* per biological replicate = 41-151 MN. (**H-I**) Percent cGAS+ MN observed by IF, 72 hours post-acute exposure to SHIELD/OHT, either as a fraction of total MN (H) or as a fraction of DNA-FISH probe+ MN (I). *N* per biological replicate = 50 MN. Statistical comparison by Student’s t-test. ns: *p* > 0.05, *: *p*<= 0.05, **: *p*<= 0.01, ***:*p*<= 0.001, ****: *p*<= 0.0001. Data are represented as mean ± SEM.

We considered whether cGAS and transcriptional machinery were competing to bind the same regions of micronuclear chromatin. In irradiated cGAS-knockout cells, we found no compensatory increase in EU staining when cGAS is lost, suggesting that cGAS does not block transcription in MN (**Fig. 2D**). Given that transcription stops when MN rupture, the occlusion of cGAS by transcription machinery would primarily apply to intact MN (Hatch et al., 2013). Yet, even when % EU+ MN is low, intact MN rarely retain cGAS, suggesting that active transcription is not blocking cGAS (**Fig. 1J, Fig. 2B**). Thus, despite a clear negative correlation between cGAS+ and EU+ MN, our data indicate this is not a simple model of competition. To more directly test the response of cGAS to MN originating from actively transcribing regions of the genome, we employed a U2OS cell-based reporter system in which DNA breaks can be created at a defined locus upstream of a transcriptional unit. Transcription at the site of DNA damage can be toggled on and off using a doxycycline (DOX)-inducible promoter (Shanbhag et al., 2010; Shanbhag and Greenberg, 2013; Tang et al., 2013). In this system, break induction led to increased MN suggesting the MN originate from the targeted locus (**Fig. 2E**); this was confirmed with DNA FISH for the targeted locus (**Fig. 2F-G**). Damage alone did not change the fraction of cGAS+ MN, but in cells where DOX-induced transcription of the locus is ongoing at the time of DNA damage, % cGAS+ MN was significantly decreased (**Fig. 2H-I**). Thus, just as is observed with acute DRB or siSRSF1 treatments, using the DOX-inducible system to target DSBs to regions of active transcription results in low cGAS-MN retention. When a locus is actively transcribing at the time of DNA damage, our data suggest that there are intrinsic factors associated with the chromatin that prevent cGAS recruitment to MN upon envelope rupture.

### Chronic DOT1L inhibition dissuades cGAS recognition of MN following IR

Actively transcribing units have a distinct chromatin architecture and histone modification profile, and attract a unique subset of remodelling and processing proteins. All of these elements could be carried into the MN produced from a DSB originating at or near a gene body, and could influence cGAS recruitment. To evaluate whether there are specific chromatin features that influence cGAS micronucleation, we screened MCF10A cells treated with a library of 48 chemical inhibitors, each designed to inhibit a specific histone reader, writer, or eraser (**Fig. 3A; Table S1**) (Scheer et al., 2019; Wu et al., 2019). After four days of inhibitor treatment, which is sufficient time to allow any target histone modifications to change within the nucleus, the cells were exposed to 10 Gy IR and allowed 72 hours to progress through the cell cycle and form MN while still in the presence of the epigenetic inhibitors. We found that cGAS recognition of IR-induced MN, which normally occurs ~80% of the time, could be significantly decreased by chronic inhibition of certain histone readers, writers, and erasers (**Fig. 3A**). Just as was seen across distinct agent exposures, the changes to % cGAS+ MN did not correlate with changes to the overall number of MN produced, further supporting the idea that cGAS recognition is an effect of the MN itself rather than the quantity of unresolved DSBs or overall MN density (**Fig. S3B**, *R*^2^ = 0.12).

**Figure 3.**
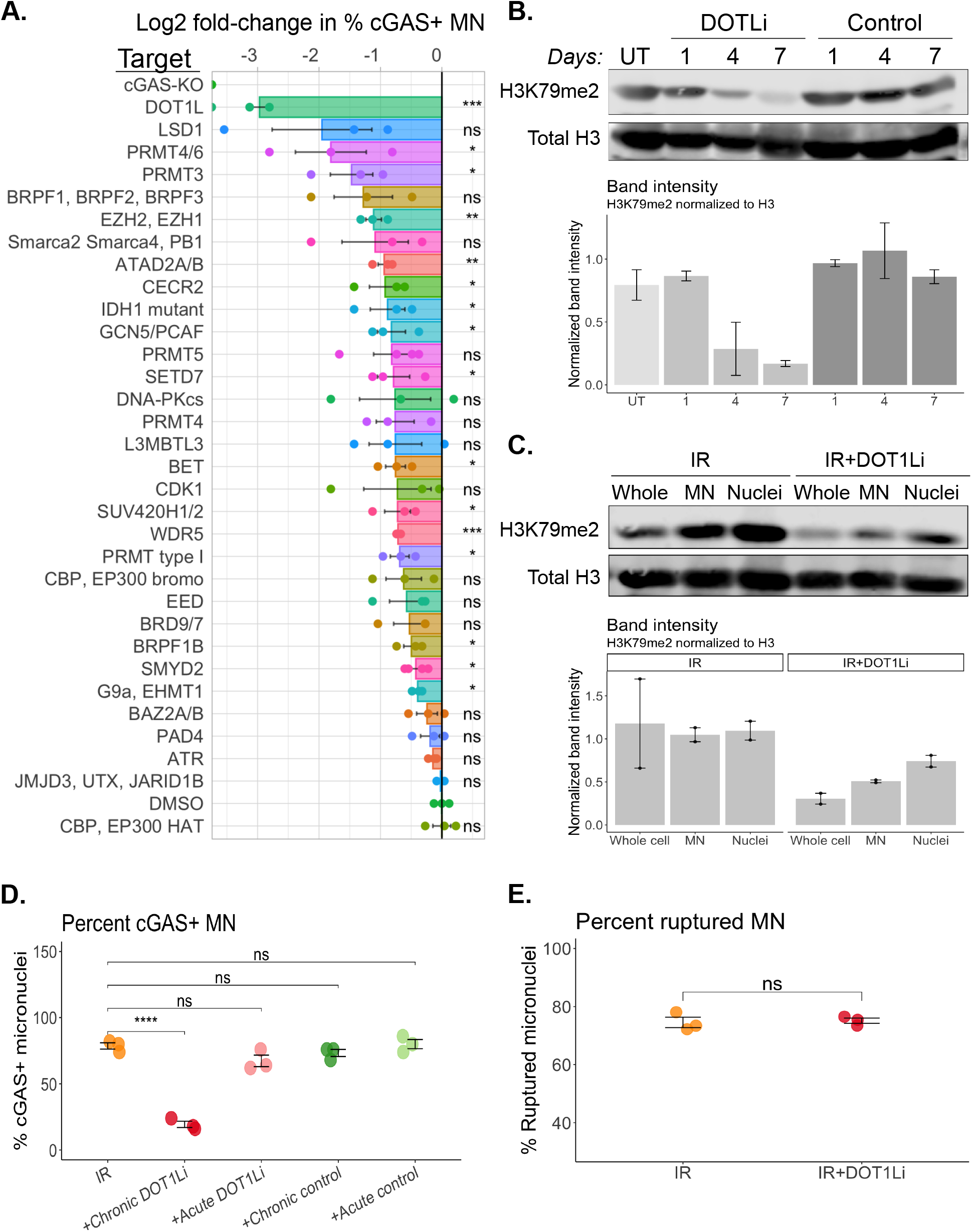
Chronic DOT1L inhibition dissuades cGAS recognition of MN following IR. (**A**) Percent cGAS+ MN observed by IF, 72 hours post-exposure to 10 Gy IR. Treatment of MCF10A cells with selective inhibitors for each of the indicated targets began four days prior to IR exposure. Data are represented as the log2 fold-change in % cGAS+ MN, as compared to cells treated with DMSO prior to IR. Statistical comparisons by Student’s t-test using IR+DMSO as the reference group. *N* per biological replicate = 50 MN. (**B**) Western blot showing H3K79me2 levels in untreated MCF10A cells, or cells treated for 1-7 days with SCG0946 (DOT1L inhibitor). Samples were acid-extracted to purify histones prior to loading. Representative of 2 replicate experiments. Intensity measurements are first normalized to blot background, then to total H3. (**C**) Western blot showing H3K79me2 levels in whole cells, purified MN, and purified nuclei, 3 days following IR exposure. IR+DOT1Li conditions were pre-treated with DOT1Li for four days prior to IR exposure. Samples were acid-extracted to purify histones prior to loading. Representative of 2 replicate experiments. Intensity measurements are first normalized to blot background, then to total H3. (**D**) Percent cGAS+ MN observed by IF, 72 hours post-exposure to 10 Gy IR. Treatment of MCF10A cells with DOT1Li or its control compound (SGC0649) began four days (chronic) or one hour (acute) prior to IR exposure. *N* per biological replicate = 50 MN. Statistical comparison by Student’s t-test. (**E**) Percent ruptured MN observed by IF, 72 hours post-exposure to 10 Gy IR. IR+DOT1Li MCF10A cells were pre-treated with DOT1Li for four days prior to IR exposure. *N* per biological replicate = 50 MN. Statistical comparison by Student’s t-test. ns: *p* > 0.05, *: *p*<= 0.05, **: *p*<= 0.01, ***: *p*<= 0.001, ****: *p* <= 0.0001. Data are represented as mean ± SEM. See also Figure S3.

The manipulation of % cGAS+ MN when cells are irradiated in the presence of certain epigenetic inhibitors suggested two non-exclusive scenarios. First, that cGAS is directly interacting with specific histone modifications on MN chromatin, and their loss prevents cGAS recognition; second, that cGAS prefers certain chromatin architectures, and the loss of specific histone modifications in the nucleus lessens the likelihood of these structures becoming sequestered into MN. To test these scenarios, we focused on the effects of SGC0946, which selectively inhibits the methyltransferase activity of DOT1L to globally erase dimethylation on lysine 79 of histone H3 (H3K79me2) (Yu et al., 2012) (**Fig. 3B-C**). Targeting DOT1L with SGC0946 had the strongest and most consistent effect on % cGAS+ MN in our screen (**Fig. 3A**). Treatment with a structurally similar control compound that does not inhibit DOT1L (SGC0649, (Yu et al., 2012)) did not alter % cGAS+ MN following IR (**Fig. 3D**). SGC0946 (hereafter referred to as DOT1Li)-induced decrease in cGAS+ MN was only observed when cells were treated with the inhibitor chronically (beginning four days prior to IR exposure) with no effect observed following acute treatment (one hour prior) (**Fig. 3D**). Thus, the reduction in cGAS+ MN is most likely an effect of nuclear erasure of H3K79me2 prior to DNA damage, rather than a consequence of inhibited DOT1L in the immediate response to damage.

To verify that chronic DOT1Li-reduced % cGAS+ MN was an effect of changes to nuclear (and by extension, micronuclear) chromatin, we ruled out alternative explanations for how DOT1Li could be affecting cGAS. Since H3K79me2 is a marker of active enhancers and promoters, we considered whether chronic DOT1Li treatment may prevent cGAS expression. By RT-qPCR and Western blot, we found that cGAS transcript and protein levels remained unchanged over 7 days of DOT1Li exposure (**Fig. S3C-D**) and the DOT1Li-driven reduction of cGAS+ MN was observed even in cells expressing a transgenic, constitutively expressed mCherry-cGAS construct (**Fig. S3E**). DOT1Li-treated cells displayed no difference in their incidence of MN envelope rupture compared to untreated, meaning variable access is not responsible for the observed difference in % cGAS recognition (**Fig. 3E**). Together, our data suggests that the DOT1Li-driven reduction in IR-induced cGAS+ MN is most likely due to loss of nuclear H3K79me2, occurring prior to DNA damage.

### cGAS binds H3K79me2, but the interaction is neither sufficient nor necessary for localization to MN chromatin

We next considered whether cGAS was interacting with specific modifications (such as H3K79me2) directly, and whether such direct interaction was necessary for cGAS localization to MN. FLAG-tagged cGAS did co-immunoprecipitate (IP) with H3K79me2, and not with H2BK120ub, H3K9ac, or H3K9me2, supporting the notion that cGAS and H3K79me2 are specific binding partners (**Fig. 4A**). Therefore, the overall reduction in cGAS+ MN under IR+DOT1Li conditions is likely influenced by the loss of a binding partner, H3K79me2, on chromatin (**Fig. 3A-D**).

**Figure 4.**
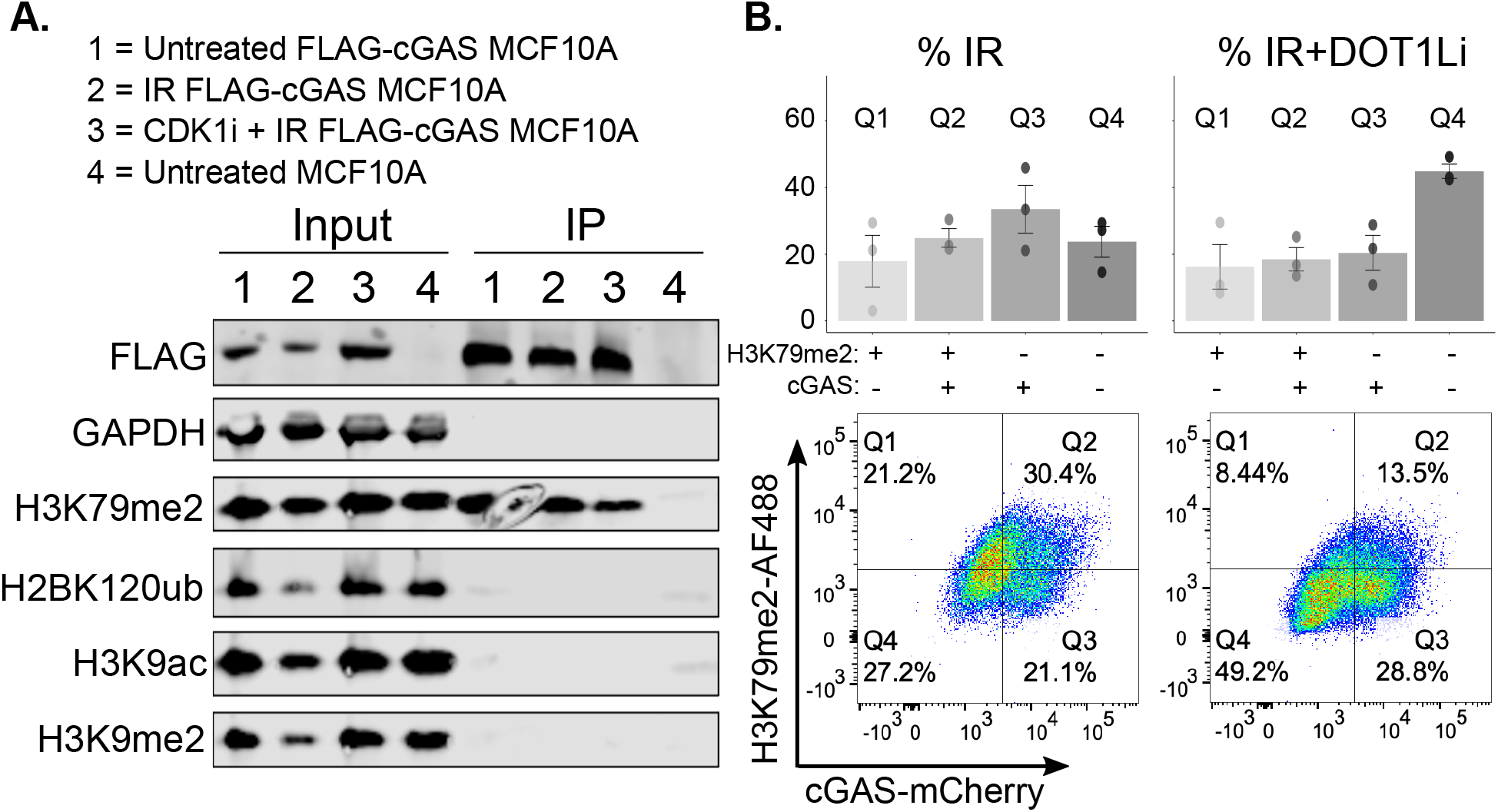
cGAS binds H3K79me2 specifically, but not exclusively. (**A**) FLAG-cGAS immunoprecipitated (IP) from MCF10A cells 72 hours following each of the four indicated treatment conditions. CDK1 inhibitor was added one hour prior to IR exposure. Representative of 2 replicate experiments. (**B**) Flow cytometry depicting the relationship between H3K79me2+ and cGAS+ MN, in MN purified from irradiated HeLa cells. IR+DOT1Li cells were pre-treated with DOT1Li for four days prior to IR exposure. Gating scheme for MN is depicted in Figure S1A. Representative of 3 replicate experiments. Data are represented as mean ± SEM. See also Figure S4.

DOT1L is the only histone methyltransferase to deposit H3K79me2, yet the inhibition of several epigenetic readers, writers, and erasers had a significant effect on cGAS localization to MN following IR (**Fig. 3A**). Having identified that H3K79me2 is positively associated with cGAS binding to MN, we wondered whether this modification was required for cGAS localization. After IR, most cGAS+ MN are also H3K79me2+, in HeLa cells evaluated by flow cytometry (**Fig 4B**). However, a substantial fraction of MN are H3K79me2+ cGAS-, or H3K79me2-cGAS+. Thus, binding to H3K79me2 appears beneficial, but neither sufficient nor necessary for cGAS retention to MN (**Fig 4B**). With DOT1Li treatment, the MN profile shifts, with most cGAS+ MN occupying the H3K79me2-fraction (**Fig. 4B**). There are likely a range of chromatin structures and states that attract and dissuade cGAS binding. We have found that these include, but are not limited to, dimethylation of H3K79 (**Fig. 3A-D, Fig. 4**) and active transcription at the time of DNA damage induction (**Fig. 2**). Chronic DOT1L inhibition does not affect the % EU+ MN, and enriching MN for active gene bodies by acute DRB treatment does not decrease the overall % H3K79me2+ MN, suggesting that these structures operate independently (**Fig. S4A-B**).

### Differential cGAS recognition of MN influences the downstream ISG cascade

cGAS recognition of MN DNA initiates a pro-inflammatory gene expression program (Harding et al., 2017; Mackenzie et al., 2017). dsDNA-bound cGAS dimerizes to produce cGAMP, and cGAMP signals through STING to initiate an inflammatory cascade comprised of dozens of genes, including ISG54 and ISG15 (Barber, 2015; Chen et al., 2020). Having observed that distinct genotoxic exposures generate MN that are differentially bound by cGAS, we assessed whether this phenomenon has consequences for the damaged cell’s ability to produce ISGs. MCF10A cells acutely exposed to individual genotoxic stressors varied in their ISG responses at 72 hours post-exposure, following a distribution that generally overlapped with the exposure’s % cGAS+ MN at the same time point (**Fig. 5A-B**). The ISG response depended on mitosis-associated MN formation in general, and on cGAS recognition of MN in particular, as cGAS-knockout or G2 arrest with a CDK1 inhibitor blocked ISG production (**Fig. 5A-B**). Consistent with our observations on cGAS recognition of MN, chronic DOT1Li treatment blunted the IR-induced ISG response (**Fig. 5C-F**). This is not due to dependence on H3K79 methylation for ISG signalling, since both cGAS and STING expression remain intact in DOT1Li-treated conditions (**Fig. S3C, Fig. S4C**) and the addition of exogenous cGAMP or herring testis (HT)-DNA can drive ISG54 and ISG15 expression in IR+DOT1Li conditions (**Fig. 5C-F**). Together, these results indicate that changes to nuclear chromatin organization (i.e. induced by DOT1Li or active transcription) occurring prior to DNA-damaging exposures can dissuade cGAS recognition of MN, and impair the cell’s ability to initiate a genotoxic stress-induced inflammatory cascade.

**Figure 5.**
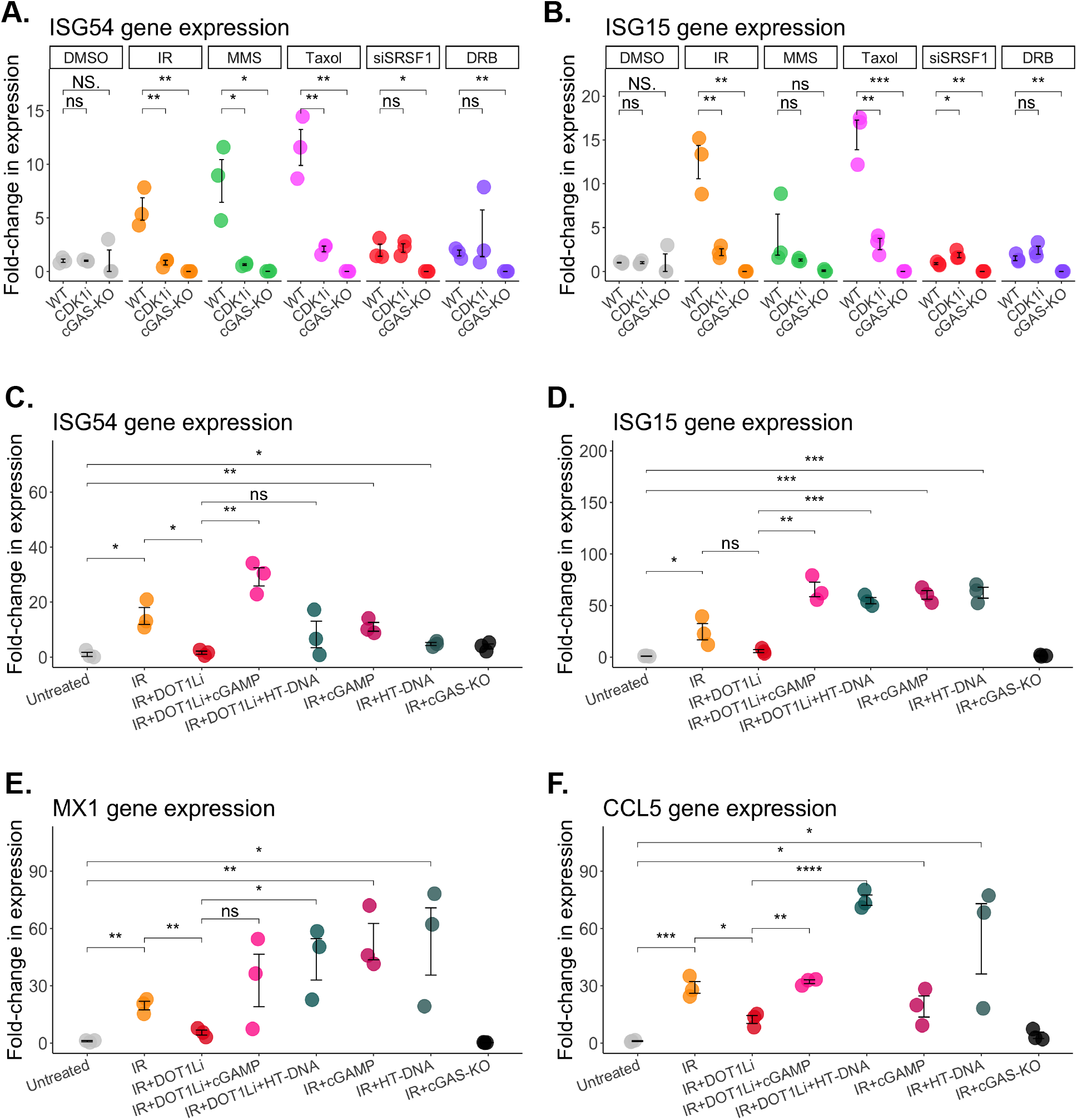
Differential cGAS recognition of MN influences the downstream ISG cascade. (**A-B**) RT-qPCR for (A) ISG54 and (B) ISG15 expression in MCF10A cells, 72 hours post-acute exposures. CDK1 inhibitor was added one hour prior to acute exposure. Fold-change in expression is represented relative to average ISG54 or ISG15 in the DMSO-treated samples. GAPDH was used as the housekeeping gene. Statistical comparison by Students t-test. (**C-F**) RT-qPCR probing for (C) ISG54 (D) ISG15 (E) MX1 and (F) CCL5 expression in MCF10A cells, 72 hours post-IR. DOT1Li treatment began four days prior to IR exposure. 100 μg/mL cGAMP or HT-DNA were added 24 hours prior to IR exposure. Fold-change in expression is represented relative to the average expression level of the untreated samples. GAPDH was used as the housekeeping gene. Statistical comparison by Students t-test. NS: p = 1, ns: *p* > 0.05, *: *p* <= 0.05, **: *p* <= 0.01, ***: *p* <= 0.001, ****: *p* <= 0.0001. Data are represented as mean ± SEM. See also Figure S4.

## DISCUSSION

We have demonstrated that distinct and common forms of DNA damage, including base alkylation and transcription stress, strongly influence the cell’s ability to produce cGAS-recognizable MN. The differential magnitude of cGAS+ MN across genotoxic agent exposures was not a function of MN envelope integrity, and instead is dictated by differences in MN DNA content, arising from the specific chromatin context of the nuclear DNA fragment that becomes MN-sequestered by each exposure. We showed that manipulating nuclear chromatin prior to DNA damage influences MN chromatin content, and this affects cGAS recruitment. In particular, depleting H3K79 dimethylation or enriching regions of active or stalled transcription for micronuclear sequestration negatively impacted MN-cGAS associations. Differential cGAS recognition of MN generated by distinct, acute genotoxic stressors translated to varying magnitudes of the cell’s cGAS-dependent ISG response. Our data contributes to the growing body of work demonstrating that micronuclear content dictates micronuclear characteristics, and expands upon the role of specific kinds of DNA damage responses in carcinogenic processes, including dsDNA-dependent inflammatory gene expression (Crasta et al., 2012; Zhang et al., 2015; Umbreit et al., 2020; Mammel et al., 2022).

When actively transcribing regions are sequestered in MN, cGAS is absent (**Fig. 2**). cGAS might be avoiding micronuclear chromatin that has come from regions of active transcription because its canonical ligand, dsDNA, is obstructed. Transcriptionally active regions contain an open DNA helix, mRNA, R-loops, polymerases, and processing enzymes, all of which could physically obstruct cGAS access to dsDNA. However, existing work has demonstrated that transcriptional machinery vacates MN upon rupture (Hatch et al., 2013). This suggests that even if MN generated from actively transcribing regions had carried obstructive machinery to the MN, MN DNA should be recognizable upon rupture. Yet, those MN still do not attract cGAS (**Fig. 1K**, siSRSF1 and DRB). We have demonstrated that cGAS localization to MN is sensitive to the presence of specific histone modifications, but the full set of structures that attract or dissuade cGAS binding is not yet known (**Fig. 3A, Fig. 4C**). Therefore, it is possible that even if transcription-stress-generated MN do contain accessible DNA, it may be structured in a way that is cGAS-averse. From recent literature describing nuclear cGAS binding, there is a precedent for cGAS exhibiting a binding preference for distinct chromatin states (Gentili et al., 2019; Michalski et al., 2020; Pathare et al., 2020; Zhao et al., 2020). Given that certain genotoxic exposures appear more likely to generate MN containing an active transcriptional unit (**Fig. 2B**), our findings have implications for the ability of a cell to signal ongoing DNA damage to the immune system via cGAS, depending on the nature of the MN-generating genotoxic insult.

cGAS directly interacts with H3K79me2, but is capable of localizing to MN even when this modification is absent (**Fig. 4**). Given this, it is interesting that the chronic inhibition of DOT1L and the loss of H3K79me2 so substantially ablates cGAS+ MN (**Fig. 3A-D**). It may be that chronic DOT1Li positions as-yet-uncharacterized cGAS-averse chromatin states within the MN, which would dissuade cGAS localization in two ways: By the loss of a binding partner (H3K79me2) and the presence of an unknown negative regulator, adding to the growing literature on cGAS binding partners and negative regulators within both nuclear and micronuclear chromatin (Michalski et al., 2020; Pathare et al., 2020; Zhao et al., 2020; Li et al., 2021; Mohr et al., 2021). Future work characterizing MN content, particularly under conditions of diverse nuclear organizational states, will continue to illuminate these relationships. These studies have potential therapeutic implications: DOT1L inhibitors are currently in clinical trials for the treatment of mixed-lineage leukemia, targeting increased expression of oncogenes under the control of H3K79me2-marked promoters (Nguyen et al., 2011). Our results suggest that DOT1L inhibitors may be useful for other types of cancers, particularly those relying on a cGAS-STING-driven inflammatory microenvironment for their persistence or metastasis (Bakhoum et al., 2021). Fully characterizing the cGAS interaction network could allow us to manage the DNA damage-induced consequences of MN formation in a targeted manner.

Interferon-stimulated gene expression is a major functional consequence of cGAS micronucleation (Harding et al., 2017; Mackenzie et al., 2017; Mohr et al., 2021). Here we have demonstrated that populations of cells exposed to different forms of acute genotoxicity exhibit variable ISG responses, following a distribution that generally overlaps with their % cGAS+ MN (**Fig. 5A-B**). cGAS micronucleation is an important event in the ISG cascade, since cGAS knockout or pre-treating cells with a CDK1 inhibitor, which prevents cells from entering mitosis and producing MN, was sufficient to block ISG production following IR, MMS, and Taxol exposure (**Fig. 5A-B**). Explicitly manipulating % cGAS+ MN after IR with DOT1Li produced corresponding changes in ISG expression (**Fig. 5C-F**). These data highlight the involvement of cGAS and MN in the ISG cascade, but there are likely many factors involved in translating MN production into a measurable ISG response. In our hands, DMSO, siSRSF1, and DRB treatments produced MN that are recognized by cGAS at non-zero frequencies, and yet they initiated no measurable ISG response (**Fig. 1C, Fig. 5A-B**). This might be explained by low MN burden under these conditions (**Fig. 1D**). However, even in cases where the total MN burden remains unchanged, such as IR+DOT1Li (**Fig. S3B**), the loss of cGAS+ MN is sufficient to abrogate ISG production (**Fig. 5C-F**). Exploring other factors involved in the cellular MN response could help explain this gap between cGAS+ MN and ISG signalling observed under conditions such as DRB or siSRSF1 treatments. Other cellular stressors can serve as co-activators of cGAS, as was recently described in the context of ongoing translation stress (Wan et al., 2021). There are inflammatory sources of self-DNA within the cell other than MN, including mitochondrial DNA (West et al., 2015; Maekawa et al., 2019; Tigano et al., 2021; Wan et al., 2021) and there are MN-binding proteins that are known negative regulators of cGAS, such as TREX1 (Mohr et al., 2021). cGAS-driven inflammatory signalling influences autoimmune progression (Uggenti et al., 2020; Zhou et al., 2021), tumour metastasis (Qing Chen, Adrienne Boire, Xin Jin, Manuel Valiente, 2016; Bakhoum et al., 2021), and the outcome of radiation therapy (Deng et al., 2014; Vanpouille-Box et al., 2017). Understanding the full scope of factors that respond to DNA damage-induced MN will clarify their role in the consequent inflammatory response.

We have linked specific genotoxic exposures to the production of MN which are variably recognized by cGAS following their envelope rupture, and which instigate differential magnitudes of inflammatory signalling. Relying on DOT1Li-driven nuclear chromatin changes and the selective induction of transcription, we have demonstrated that cGAS recognition of MN is dictated by the status of the enclosed MN DNA, dependent on the existing nuclear chromatin status at the damage site. These results add to a growing consensus that cGAS localization and activity is determined by chromatin status, and promotes a more complete understanding of MN activity when generated following distinct forms of DNA damage (Sepaniac et al., 2021; Mammel et al., 2022).

## ACKNOWLEDGMENTS

We thank all members of the Harding lab for critical comments on the manuscript; R. Greenberg for U2OS reporter cells; S. Ackloo for assistance with the Structural Genomics Consortium (SGC) epigenetic inhibitor collection. This work in the Harding lab was supported by the Canadian Institutes for Health Research (CIHR), The National Science and Engineering Research Council of Canada, the JP Bickell Foundation, the Princess Margaret Cancer Centre, the Princess Margaret Cancer Foundation and Ontario Ministry of Health. KMM has received support from the Ontario Graduate Scholarship, CIHR and the University of Toronto Endowed Awards; MMT has received support from the Strategic Training in Transdisciplinary Radiation Sciences (STARS21) Program. The SGC is a registered charity (no: 1097737) that receives funds from Bayer AG, Boehringer Ingelheim, Bristol Myers Squibb, Genentech, Genome Canada through Ontario Genomics Institute [OGI-196], EU/EFPIA/OICR/McGill/KTH/Diamond Innovative Medicines Initiative 2 Joint Undertaking [EUbOPEN grant 875510], Janssen, Merck KGaA (aka EMD in Canada and US), Pfizer and Takeda.

## AUTHOR CONTRIBUTIONS

KMM and SMH designed the experiments and wrote the manuscript with input from all authors. SNP performed the DNA FISH experiments presented in Fig. 2F-I and the IF assessments presented in Fig. S1B, and created mCherry-cGAS/GFP-NLS-expressing MCF10A cells used for live cell imaging experiments. MMT performed the experiment presented in Fig. S3A. KMM performed and analyzed all other experiments. CA provided the inhibitor library presented in Fig. 3A, guided the design of experiments, and edited the manuscript.

## METHODS

### EXPERIMENTAL MODEL AND SUBJECT DETAILS

MCF10A cells were cultured in 1:1 mixture of F12:DMEM media supplemented with 5% horse serum (Wisent Bioproducts cat #098150), 20 ng/ml human EGF (Cedarlane Labs cat #AF-100-15), 0.5 mg/ml hydrocortisone (Sigma-Aldrich cat #H0888), 100 ng/ml cholera toxin (Sigma Aldrich cat #C8052) and 10 mg/ml recombinant human insulin (SAFC cat #91077C). HeLa, U2OS, and HEK293T cells were grown in DMEM supplemented with 10% FBS. All media was supplemented with 1% penicillin-streptomycin.

### METHOD DETAILS

#### Acute treatments with varied genotoxic stressors

Treatments were performed on cultured cells. Doses and exposure times were selected based on existing literature and preliminary experiments to induce DNA lesions and maximize MN formation, while minimizing cell death and cell cycle arrest within the time frame of the assay (72 hours). Doses and exposure times used for each agent are as follows: ionizing radiation (IR, Cs-137), 10 Gy (Harding et al., 2017); methylmethanesulfonate (MMS, Sigma-Aldrich cat #12995), 0.5 μM for 4 hours (Lutz et al., 2005); ultraviolet radiation (UV, 254 nm), 15 J/m^2^ (Oksenych et al., 2013); paclitaxel (Taxol, Selleck Chemicals cat #S1150), 10 nM for 4 hours (Xu et al., 2011); hydroxyurea (HU, Sigma-Aldrich cat #148627), 2 mM for 24 hours (Utani et al., 2010; Xu et al., 2011); cisplatin (Millipore cat #232120), 0.5 μg/mL for 24 hours (Zhao et al., no date); siSRSF1 (Thermo Fisher Scientific, cat #4427037, ID #s12725), 2 nM for 24 hours (Cohen S, Guenolé A, Marnef A, Clouaire T, Puget N, Rocher V, Arnould C, Aguirrebengoa M, Genais M, Vernekar D, Mourad R. Borde, 2020); mitomycin C (MMC, Sigma cat #M4287), 0.5 μg/mL for 4 hours (Bakhoum et al., 2018; Sommer, Buraczewska and Kruszewski, 2020); 5,6-dichloro-1-beta-ribo-furanosyl benzimidazole (DRB, Sigma-Aldrich cat #D19116), 100 μM for 4 hours (Cohen S, Guenolé A, Marnef A, Clouaire T, Puget N, Rocher V, Arnould C, Aguirrebengoa M, Genais M, Vernekar D, Mourad R. Borde, 2020). At the end of the designated exposure time, cells were washed in PBS, replaced with fresh media, and allowed to cycle freely for 72 hours to produce MN before proceeding to downstream analysis.

#### Immunofluorescent microscopy

For immunofluorescent microscopy (IF), cells were seeded onto glass coverslips 24 hours prior to treatments with acute genotoxic stressors. After treatment, cells on coverslips were washed twice with PBS + 0.1% Tween-20 (PBS-T). Cells were fixed for 10 minutes on ice with 3% paraformaldehyde (PFA) + 2% sucrose in PBS. Cells were washed twice in PBS-T, then permeabilized for 10 minutes at room temperature with 0.5% NP-40 solution in PBS. Cells were washed twice in PBS-T, incubated in blocking solution (3% bovine serum albumin [BSA] in PBS-T) for 10 minutes at room temperature. Cells were stained with the appropriate primary antibodies for a given experiment, overnight at 4 °C. The following dilutions were used for primary antibodies: 1:1000 cGAS (Cell Signalling Technologies cat #15102S); 1:2000 CENP-A (Abcam cat #ab13939); 1:1000 γH2AX (Millipore cat #05-636), 1:250 CHMP7 (Proteintech cat #16424-1-AP). Cells were washed four times in PBS-T, then incubated in fluorescently labelled secondary antibodies for 1 hour at room temperature. The following dilutions were used for secondary antibodies: 1:500 AlexaFluor 568 Goat anti-rabbit IgG secondary (Invitrogen cat #A11036); 1:500 AlexaFluor 488 Goat anti-mouse IgG secondary (Invitrogen cat #A11001); 1:500 AlexaFluor 568 Goat anti-mouse IgG secondary (Invitrogen cat #A11004); 1:500 AlexaFluor 488 Goat anti-rabbit IgG secondary (Invitrogen cat #A11034); Cells were washed four times in PBS-T, then inverted onto a drop of Prolong Glass Antifade Mountant with NucBlue (DAPI) stain (Thermo Fisher Scientific, cat #P36981) on a microscope slide. Slides were imaged on an Olympus immunofluorescent microscope, using CellSens Dimensions software. All IF images presented here were taken at 60X objective on an oil lens.

#### Live cell imaging

Live cell imaging of MCF10A cells stably expressing GFP-tagged nuclear localization signal (GFP-NLS), mCherry-tagged cGAS (mCherry-cGAS), or mCherry-tagged histone 2B (mCherry-H2B) were performed using an Incucyte SX5 (Sartorius Imaging). Images were taken once every 60 minutes to acquire videos of cells monitored over a 72-hour period.

#### Viral transfection and selection of stable cell lines

To prepare MCF10A and HeLa cells stably expressing GFP-NLS, FLAG-cGAS, mCherry-cGAS, and/or mCherry-H2B, lentiviral transfection followed by fluorescence-activated cell sorting (FACS) or antibiotic selection was performed. pTRIP-SFFV-EGFP-NLS (Addgene ID #86677), pOZ-cGAS (puromycin-resistant), pLXV-mCherry-cGAS (puromycin-resistant), and/or pLenti6-mCherry-H2B (Addgene ID #89766, blasticidin-resistant) were transfected into HEK293T cells as follows, according to LipoD293 reagent instructions (FroggaBio cat #SL100668.1): Lentiviral packaging plasmids pMD2.g (Addgene #12269), pRSV-Rev (Addgene #12253), and PMDLg/pRRE (Addgene #12251) were combined in serum-free DMEM, using 0.5 μg of each plasmid for 125 μL of DMEM. 1 μg of target plasmid was added to this mixture per 125 μL DMEM. In a separate tube, 7.5 μL of LipoD293 was added per 125 μL DMEM. The two mixtures were combined and allowed to incubate at room temperature for 30 minutes, then added dropwise to HEK293T cells cultured in DMEM supplemented with 10% FBS and 1% penicillinstreptomycin. Cell supernatants containing lentiviruses were mixed 1:1 with target cell media and supplemented with 4 mg/ml polybrene (Sigma-Aldrich cat #107689). Successfully transduced cells were selected using puromycin (Cedarlane cat #13884-50), blasticidin (Cedarlane cat #14499-25), or in the case of GFP-NLS-infected cells, FACS sorting. pOZ-cGAS and pLXV-mCherry-cGAS plasmids were cloned as previously described (Harding et al., 2017).

#### Micronuclei purification

Micronuclei were partially purified prior to flow cytometric analysis according to a previously described protocol (Mohr et al., 2021). 500 mL suspension-cultured HeLa cells pre-treated with acute genotoxic stressors were harvested and washed twice in DMEM without serum. Cells were resuspended in DMEM without serum, supplemented with cytochalasin B (BioShop Canada cat #CYT444.25) at 10 μg/ml, and incubated at 37 °C for 30 minutes. Cells were centrifuged at 250 x g for 5 minutes and the cell pellet was resuspended in ice-cold lysis buffer (10 mM Tris-HCl pH 8.0, 0.32 mM sucrose, 2 mM magnesium acetate, 3 mM calcium chloride, 0.1 mM EDTA, 0.1% NP-40, pH adjusted to 8.5) freshly supplemented with 10 μg/mL cytochalasin B, 0.15 mM spermine, 0.75 mM spermidine, 1 mM dithiothreitol (DTT) and 200 mM sodium butyrate (Sigma). Resuspended cells were homogenized by douncing, using 15 strokes with a loose-fitting pestle. Cell lysates were then mixed with an equal volume of ice-cold 1.8 M sucrose buffer (10 mM Tris-HCl pH 8.0, 1.8 M sucrose, 5 mM magnesium acetate, 0.1 mM EDTA, 0.3% BSA, pH adjusted to 8.0) freshly supplemented with spermine, spermidine, DTT, and sodium butyrate. 10 mL of this mixture was layered onto a two-layer sucrose gradient, prepared by adding 20 mL of 1.8 M sucrose buffer on top of 15 mL of 1.6 M sucrose buffer in a 50 mL conical tube. The layers are added to each slowly, such that the middle 1.8 M buffer layer does not mix with the lower 1.6 M buffer layer, and the upper 1:1 mixture of lysed cells + 1.8 M does not mix with the 1.8 M buffer layer. The gradient was centrifuged for 1000 g for 20 minutes at 4 °C, during which time the nuclei and micronuclei contained within the lysed mixture will distribute within the gradient according to size. Fractions can be generally collected as follows: Upper 3 mL contains debris and is discarded, next 6-12 mL contains MN with minimal contaminating nuclei and is collected for flow cytometry, final 35 mL contains primary nuclei and is not used. At this point, the MN partially purified from treated conditions are processed for flow cytometry. A control sample will continue through full purification, to get rid of minimal contaminating nuclei in the MN fractions and serve as a reliable size gate for flow cytometry (**Figure S1A**). Full purification steps are adapted from a published procedure (Shimizu, Kanda and Wahl, 1996) and are as follows: MN- and contaminating nuclei-containing fractions were layered on top of a 5 mL cushion of 1.2 M sucrose buffer, in a 50 mL ultracentrifuge tube (Beckman Coulter cat #344058). MN were pelleted by ultracentrifugation in an Optima XPN-80 ultracentrifuge (Beckman Coulter) at 35,000 g for 90 minutes in a SW41 rotor, at 4 °C. The supernatant was discarded and the pellet was resuspended in 500 μL of 1.0 M sucrose buffer. The pellet was layered on top of an 11 mL linear 1.0 – 1.8M sucrose buffer gradient, prepared in a 15 mL Falcon tube as follows: Add 2.75 mL of 1.8 M sucrose buffer to the bottom of the tube. Freeze on liquid nitrogen or by brief storage at −80 °C. Layer 2.75 mL of a 2:1 mixture of 1.8 M: 1.0 M sucrose buffer onto the frozen layer. Freeze new layer as before. Layer 2.75 mL of a 1: 2 mixture of 1.8 M: 1.0 M sucrose buffer onto the frozen layer. Freeze new layer as before. Layer 2.75 mL of 1.0 M sucrose buffer onto the frozen layer. Freeze new layer as before. Thaw entire gradient upright at 4 °C when ready for use. MN pellet was centrifuged through the 11 mL linear gradient at 530 g for 15 minutes at 4 °C. Fractions can be generally collected as follows: Upper 1-4 mL contains pure MN is collected for flow cytometry, final 8 mL contains primary nuclei and is discarded.

#### Flow cytometry on purified MN

MN were purified as described above and pelleted by centrifugation at 1500 g x 3 minutes. MN were washed twice in 2% FBS in PBS, then resuspended in 400 μL 1% PFA in PBS for fixing, incubating for 30 minutes at 4 °C. MN were washed twice in 2% FBS in PBS, then resuspended in 400 μL 0.5% NP-40 in PBS for permeabilization, incubating for 30 minutes at 4 °C. MN were washed twice with staining buffer (0.3% BSA, 0.5% NP-40, and 1% sodium azide in PBS), then resuspended in appropriate primary antibody diluted 1:500 in staining buffer. Primary antibodies used for flow cytometry are: anti-H3K79ME2 (Abcam cat #ab3594). MN were incubated in primary antibody for 30 minutes at 4 °C, vortexing briefly every 10 minutes to prevent MN from settling. MN were washed twice in staining buffer, then resuspended in appropriate secondary antibody diluted 1:500 in staining buffer. Secondary antibodies used for flow cytometry are: AlexaFluor 568 Goat anti-rabbit IgG secondary (Invitrogen cat #A11036); AlexaFluor 488 Goat anti-rabbit IgG secondary (Invitrogen cat #A11034). MN were incubated in secondary antibody for 30 minutes at 4 °C, vortexing briefly every 10 minutes to prevent MN from settling. MN were washed twice in 2% FBS in PBS, then resuspended in 2% FBS in PBS to a final concentration of 1 – 10 E6 MN/mL. MN were filtered through the 35 μm mesh cap of 5 mL polystyrene roundbottom FACS tubes (Corning, cat #352235), and 50 μg/mL DAPI (Life Technologies cat #D1306) was added to the filtered samples immediately prior to conducting flow cytometry. Flow cytometry was performed on an LSRII OICR analyzer at the SickKids-UHN Flow Cytometry Core Facility.

#### Western blotting

Cells were harvested by scraping and pelleted by centrifugation at 500 g for 5 minutes. Cell pellets were resuspended in a lysis buffer composed of RIPA buffer freshly supplemented with 1 mM benzamidine HCl, 1 μg/ml antipain, 5 μg/ml aprotinin, 1 μg/ml leupeptin, 0.5 mM phenylmethylsulfonyl fluoride (PMSF), 1 mM DTT, 2 mM NaOV_4_, 10 mM NaF, 2 mM imidazole, 1.15 mM sodium molybdate, and 4 mM sodium tartrate (Sigma). Lysis was conducted on ice for 20 minutes, vortexing cells briefly after 10 minutes to prevent them from settling. Lysate was centrifuged at 18,000 g for 15 minutes at 4 °C, and the supernatant was transferred to a clean microcentrifuge tube. Protein amount was measured by Bradford assay, and 20 μg of protein was diluted to 20 μL in ddH_2_O, then combined with 5 μL of 5X sample buffer (50 mM Tris, 10% glycerol, 2% SDS, 0.01% bromophenol blue, 2.5% β-mercaptoethanol), denatured at 95 °C for ten minutes, and resolved on a Bolt™ 4-12% Bis-Tris Plus Gel (Thermo Fisher Scientific cat #NW04127BOX). Proteins were transferred from the gel to a nitrocellulose membrane (LI-COR Biosciences cat #926-31092). Membranes were blocked in 5% milk in PBS-T for 1 hour and incubated with primary antibody overnight at 4 ^°C^. Membranes were then washed 4 times in PBS-T and incubated for 1 hour at room temperature with secondary antibodies diluted 1:20,000. The following secondary antibodies were used for Western blots: Donkey Anti-Rabbit IgG AlexaFluor 790 (Jackson ImmunoResearch Labs cat #711-655-152), Donkey Anti-Mouse IgG AlexaFluor 680 (Jackson ImmunoResearch Labs cat #711-655-152). Following four washes in PBS-T, membranes were imaged using a LIC-COR.

#### Quantification of Western blot bands

For the quantifications presented in Fig, 3B-C, band intensities were measured using the region of interest (ROI) and intensity measurement tools in FIJI (ImageJ). The same ROI was used to measure the mean intensity of each band in the presented blots, as well as the intensity of the blot background. Mean H3K79me2 and total H3 band intensities were first normalized to their own blot background. H3K79me2 intensities were normalized to the lane’s corresponding total H3 band intensity for graphical presentation.

#### Gene expression analysis

Total RNA was isolated from cells using TriZol reagent (Life Technologies cat #15596018) according to the manufacturer’s instructions. cDNA was generated from 1000 ng of RNA using the iScript cDNA synthesis kit (Bio-Rad cat #1708891) according to the manufacturer’s instructions. qPCR was performed with gene-specific primers (See Key Resources Table) using SSO Advanced Universal SYBR Green Supermix (Bio-Rad cat #1725274) on a BioRad C1000Touch Thermocycle. Relative transcription levels were calculated by normalizing to GAPDH expression.

#### Immunoprecipitation

MCF10A or HeLa cells expressing FLAG-tagged cGAS were harvested by scraping and pelleted by centrifugation at 3000 g for 10 minutes at 4 °C. The cell pellet was resuspended in NET-N lysis buffer (100 nM NaCl, 50 mM Tris-HCl pH 8.0, 2 mM EDTA, 0.5% NP-40, and 10 nM MgCl_2_) freshly supplemented with 10 mM NaF, 2 mM imidazole, 1.15 mM sodium molybdate, and 4 mM sodium tartrate, 200 mM sodium butyrate, 10 units/mL benzonase, 1mM DTT, 0.5 mM PMSF, 1 mM benzamidine HCl, 1 μg/ml antipain, 5 μg/ml aprotinin, and 1 μg/ml leupeptin (Sigma). Cells were left in lysis buffer on ice for 30 minutes, vortexing briefly every 10 minutes to prevent the cells from settling. Lysate was centrifuged at 18,000 g for 15 minutes at 4 °C, and the supernatant was transferred to a clean microcentrifuge tube. Protein amount was measured with a Bradford assay. 10 μg of protein was kept as input, and 500 – 1000 μg of protein was taken for immunoprecipitation (IP). Protein for IP was diluted in NET-N buffer to a total volume of 500 – 1000 μL. FLAG-M2 beads (Sigma-Aldrich cat #A2220) were washed once in PBS and once in NET-N buffer, centrifuging 3000 rpm for 3 minutes at 4 °C to pellet the beads. Equilibrated FLAG-M2 beads were resuspended in NET-N buffer, and 10 μL beads were added to 1000 μL of IP protein solution. The microcentrifuge tubes containing the IP protein extract + FLAG-M2 beads were left at constant rotation overnight at 4 °C. The beads were pelleted by centrifugation at 3000 rpm for 3 minutes at 4 °C, and the supernatant was saved as flow-through. The beads were washed four times in 1 mL NET-N buffer, centrifuging at 3000 rpm for 3 minutes at 4 °C. The immunoprecipitate was then eluted from the beads by competition with FLAG peptide (Sigma-Aldrich cat #F47799) as follows: Pelleted beads were resuspended with 20 μL of 0.2 mg/mL FLAG peptide per 10 μL beads and left to incubate for 1 hour at 4 °C. The mixture was redistributed by pipetting every 15 minutes. At the end of the incubation, beads were pelleted by centrifugation at 3000 rpm for 3 minutes at 4 °C, and the supernatant containing eluted FLAG-cGAS was moved to a fresh microcentrifuge tube. At this point, protein samples were used for Western blotting as described above.

#### 5-ethynyl uridine (EU) staining

For IF visualization of active transcription, EU staining combined with click chemistry (Invitrogen cat #C10269) was performed. Staining was carried out according to click chemistry kit manufacturer instructions. In brief: 0.5 mM EU was added to cultured cells, one hour prior to fixation for IF. Cells were fixed and permeabilized as described for immunofluorescent microscopy. Cells were washed twice in PBS-T, then incubated with the click chemistry reaction mixture containing an AlexaFluor 488 reactive azide (Thermo Fisher Scientific, cat #A10266) for 30 minutes at room temperature. Cells were washed twice in PBS-T, then processed as described for immunofluorescent microscopy, beginning from blocking with 3% BSA in PBS-T.

#### Doxycycline-inducible transcription and directed DNA damage in U2OS cells

U2OS cells stably expressing FokI endonuclease and the 256 construct as previously described were obtained from Dr. Roger Greenberg’s lab (Shanbhag et al., 2010; Tang et al., 2013). Treatment with 4-hydroxytamoxifen (4-OHT, Sigma-Aldrich cat #H7904) and Shield-1 (Cheminpharma cat #CIP-S1-0005) stabilizes the endonuclease and allows its nuclear transduction, where it will induce DSBs at the targeted fusion protein (Shanbhag et al., 2010; Shanbhag and Greenberg, 2013). 256-expressing U2OS cells were seeded on glass coverslips 24 hours prior to treatment with 2 μg/mL doxycycline (DOX, Sigma-Aldrich cat #D3072). DOX was left overnight to drive transcription at the reporter locus. Shield-1 and 4-OHT were then added to the cells at 1 μM and 2 μM, respectively. 6 hours later, SHIELD/OHT were washed off the cultured cells, and DOX was replaced. Cells were left for 72 hours to cycle freely and generate micronuclei, before either evaluating % cGAS+ MN by IF or verifying MN content with DNA FISH.

#### DNA fluorescence *in situ* hybridization (FISH)

Cells were seeded on glass coverslips 24 hours prior to beginning FISH protocol. Cells on coverslips were washed three times in PBS, then fixed for 10 minutes at 4 °C with 2% paraformaldehyde in PBS. Cells were washed three times in PBS, then permeabilized for 10 minutes at 4 °C with 0.5% NP-40 solution. Cells were washed three times in PBS, then denatured in 50% formamide/2X sodium saline citrate (SSC) at 75 °C for 30 minutes. Cells were washed three times in PBS, then inverted onto a 5 μL droplet of hybridization buffer on a glass microscope slide. Hybridization buffer is composed of 50% formamide, 50% dextran, 20% SDS, 20X SSC, and 1 μL DNA FISH probe per 5 μL hybridization buffer. Coverslips were sealed with CytoBond (SciGene cat #2020-00-2) and the slide was incubated at 75 °C for 10 minutes, then transferred to a humidified chamber to incubate at 37 °C overnight. CytoBond was then removed, and coverslips were lifted from the slide and submerged in 2X SSC. Cells on coverslips were rinsed once in 2X SSC, then washed three times with 50% formamide/2X SSC, incubating the coverslips at 37 °C for 5 minutes between each wash. Cells were rinsed once in PBS-T, then incubated in 3% BSA in PBS-T (blocking buffer) for 30 minutes at room temperature. After blocking, cells were incubated with an anti-DIG FITC-tagged probe (Millipore cat #1120774191) diluted 1:200 in blocking buffer. Cells were incubated with the probe for 1 hour at room temperature, then washed three times in PBS-T. Coverslips were then inverted onto a drop of Prolong Glass Antifade Mountant with NucBlue (DAPI) stain (Thermo Fisher Scientific, cat #P36981) on a microscope slide. Slides were imaged as described above for immunofluorescent microscopy.

#### Epigenetic inhibitor screen

The screen presented in Figure 3A was performed using a library of chemical inhibitors, each designed to inhibit a specific histone reader, writer, or eraser (Scheer et al., 2019; Wu et al., 2019). The screen takes place in a 96-well plate containing 1000X stock solutions of each inhibitor, with concentrations chosen according to the screen manufacturer’s specifications (**Table S1**). MCF10A cells were seeded in 96-well plates in 198 μL of growth media, 24 hours prior to the start of the screen. They were seeded in a Corning high content imaging plate (cat #CLS4580), allowing for IF microscopy to take place directly within the wells. 2 μL of each 1000X inhibitor were added to the wells containing MCF10A cells, diluting to 1X. Cells were treated with the library for four days, as this is indicated by the screen’s specifications to be sufficient time for all targeted histone modifications to be affected. After four days, the plate was exposed to 10 Gy IR, and left for another 72 hours to produce MN while still in the presence of the epigenetic inhibitors. 72 hours post-IR exposure, the inhibitors were washed off. Fixation, permeabilization, blocking, and primary/secondary antibody staining were conducted as described above for IF microscopy, but on cells at the bottom of the imaging plates rather than on glass coverslips. After antibody staining, wells were filled with 150 μL of 300 nM DAPI (Life Technologies cat #D1306) in PBS, and imaged on an Olympus immunofluorescent microscope as described for IF microscopy.

#### Histone acid extraction

For the immunoblots presented in Figure 3B-C, histones were acid-extracted from lysates prior to loading, following a previously described protocol (Sidoli et al., 2016). Treated cells were harvested by scraping and pelleted by centrifugation at 500 g for 5 minutes. Cell pellets were resuspended in 200 μL nuclear isolation buffer (NIB; 15 mM Tris-HCl, 60 mM KCl, 15 mM NaCl, 5 mM MgCl_2_, 1 mM CaCl_2_, 250 mM sucrose, pH adjusted to 7.5) freshly supplemented with 0.5 mM DTT, 1 mM PMSF, 10 mM NaF, 2 mM imidazole, 1.15 mM sodium molybdate, 4 mM sodium tartrate, and 200 mM sodium butyrate (Sigma). Cells were centrifuged at 1000 g for 5 minutes at 4 °C, and the supernatant was discarded. Cell pellet was resuspended in freshly supplemented NIB + 0.2% NP-40 for lysis, and incubated on ice for 15 minutes. Cells were vortexed briefly after five minutes, to prevent them from settling. After lysis, cells were pelleted by centrifugation at 1000 g for 5 minutes at 4 °C, and the supernatant was discarded. Cells were washed twice in supplemented NIB. After the second wash, cell pellet was resuspended in 400 μL chilled 0.2 M H_2_SO_4_, and incubated for three hours at constant rotation. The extraction was then centrifuged at 11,000 g for 5 minutes at 4 °C, and the supernatant was transferred to a fresh microcentrifuge tube. 132 μL of 100% tricholoracetic acid (Sigma-Aldrich cat #T9159) was added dropwise to the supernatant, to a final concentration of 33%. The tubes were mixed by inverting, then incubated at 4 °C overnight. Cells were then pelleted by centrifugation at 16,000 g for 10 minutes at 4 °C, and the supernatant was discarded carefully, without scraping the sides or the bottom of the tube. The sides and bottom of the tube were rinsed with 1 mL of ice-cold 0.1% HCl in acetone, centrifuging 16,000 g for 10 minutes at 4 °C and carefully discarding the supernatant without scraping the sides or the bottom of the tube. The HCl wash was repeated once, then when the supernatant was carefully discarded the tubes were left open at room temperature for 20 minutes to dry. The histones coating the sides and bottom of the tube were resuspended in 20 – 40 μL of ddH2O, and at this point were used for Western blotting as described above.

#### Statistical analysis

Information regarding biological replicates, sample size, and statistical testing is supplied in the Figure Legends.

---

**Table S1.** Structural Genomics Consortium epigenetic inhibitor library. Information on target protein, inhibitor compound, recommended dose and exposure times for the library used to generate the data presented in Figure 3A, as set out by the Structural Genomics Consortium.

| Compound | Target | Recommended concentration (μM) | Minimum time required for full reduction in in-cell biomarker activity | PMID |
| --- | --- | --- | --- | --- |
| GSK8814 | ATAD2A/B | 1 | 1h | 27530368 |
| BAY-850 | ATAD2A | 3 | 1h | 29043777 |
| GSK2801 | BAZ2A/B | 3 | 1h | 25799074 |
| BAZ2-ICR | BAZ2A/B | 1 | 1h | 25719566 |
| JQ1 | BET | 0.2 | 5h | 20871596 |
| BI-9564 | BRD9/7 | 1 | 1h | 26914985 |
| TP-472 | BRD9/7 | 1 | 18h | 26914985 |
| I-BRD9 | BRD9 | 1 | 18h | 25856009 |
| NI-57 | BRPF1, BRPF2, BRPF3 | 1 | 1h | 28714688 |
| PFI-4 | BRPF1B | 1 | 1h | 26139243 |
| GSK6853 | BRPF1B | 1 | 1 hr | 27326325 |
| BAY-299 | BRPF2(1)/TAF1(2) | 1 | 1 hr | 28402630 |
| SGCCBP30 | CBP, EP300 bromo | 1 | 1h | 24946055 |
| ICBP112 | CBP, EP300 bromo | 3 | 1h | 26552700 |
| A-485 | CBP, EP300 HAT | 0.8 | 3 hr | 28953875 |
| NVS-CECR2-1 | CECR2 | 1 | 1h | 28402630 |
| TP-238 | CECR2/FALZ | 0.3 |  | 28402630 |
| GSK4027 | GCN5/PCAF | 1 | 18h | 28002667 |
| L-Moses | GCN5/PCAF | 5 | 18h | 27966810 |
| GSK J4 | JMJD3, UTX, JARID1B | 5 | 1d | 22842901 |
| GSKLSD1 | LSD1 | 1 | 1-2d | 26175415 |
| GSK484 | PAD4 | 10 | 3h | 26436839 |
| PFI-3 | Smarca2 Smarca4, PB1 | 1 | 1h | 26139243 |
| UNC1215 | L3MBTL3 | 1 | 3d | 23292653 |
| Vinspinin | Spindlin (Tudor like 1 and 2 domain) | 0.3 |  | 31260300 |
| BI-9321 | NSD3 PWWP1 | 10 | 1d | 31285596 |
| A-395 | EED | 1 | 3d | 28135237 |
| SGC0946 | DOT1L | 1 | 4-14d | 23250418 |
| UNC1999 | EZH2, EZH1 | 3 | 3d | 23614352 |
| GSK343 | EZH2 | 3 | 3d | 24900432 |
| UNC0642 | G9a, EHMT1 | 1 | 3d | 24102134 |
| A-366 | G9a, EHMT1 | 1 | 3d | 24900801 |
| MRK-740 | PRDM9 | 3 | 1d | 31848333 |
| MS023 | PRMT type I | 0.1 - PRMT1; 500 - PRMT6 | 20 h (PRMT6), 2 days (PRMT1) | 26598975 |
| MS049 | PRMT4/6 | 5 | 20h | 31657716 |
| SGC 707 | PRMT3 | 1 | 1d | 25728001 |
| TP-064 | PRMT4 | 1 | 3d | 29712619 |
| SKI-73 | PRMT4 | 1 | 3d | 31657716 |
| GSK591 | PRMT5 | 1 | 4d | 26985292 |
| LLY-283 | PRMT5 | 0.5 | q | 30034588 |
| SGC3027 | PRMT7 | 3 | 2d | 32409666 |
| PFI-2 | SETD7 | 1 | 1d | 25136132 |
| BAY598 | SMYD2 | 1 | 1d | 27075367 |
| PFI-5 | SMYD2 | 3 | 1d | 31415173 |
| BAY-6035 | SMYD3 | 1 | 1d | 34154424 |
| A-196 | SUV420H1/2 | 1 | 1-2d | 28114273 |
| OICR-9429 | WDR5 | 3 | 1d | 26167872 |
| GSK864 | IDH1 mutant | 0.3 | 2d | 25622091 |

**Figure S1.**
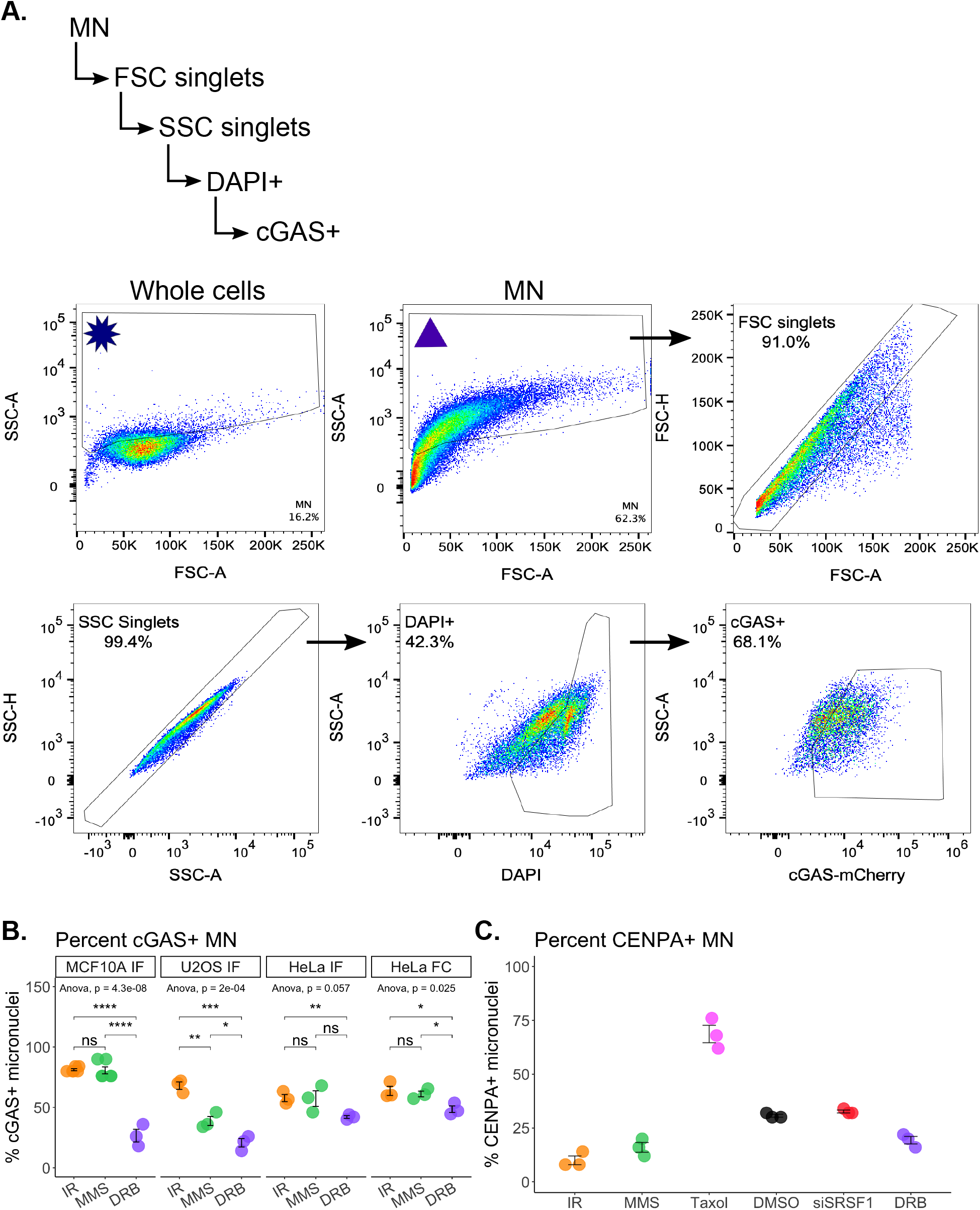
cGAS recognition of MN in MCF10A, U2OS, and HeLa cells. (**A**) Gating scheme used to evaluate cGAS+ MN by flow cytometry. MN are partially purified from experimental samples according to the protocol described in Methods and in (Mohr et al., 2021). Asterisk depicts whole cells used to set this initial size gate; events higher on the SSC axis than whole cells are considered MN. Using the size gate set by whole cells, gating of partially purified MN begins with the triangle, and proceeds according to the arrows. Gates for fluorophores including DAPI and cGAS were set according to the corresponding fluorescence-minus-one (FMO) sample. (**B**) Percent cGAS+ MN observed by IF or flow cytometry, 72 hours post-acute exposure. For IF experiments, *n* per biological replicate = 50 MN. For flow cytometry, *n* per biological replicate = all MN partially purified from a treated culture, evaluated as depicted in Figure S1A. Statistical comparisons by Students t-test and two-way ANOVA. (**C**) Percent CENPA+ MN observed by IF, 72 hours post-acute exposure. *N* per biological replicate = 50 MN. ns: *p* > 0.05, *: *p* <= 0.05, **: *p* <= 0.01, ***: *p* <= 0.001, ****: *p* <= 0.0001. Data are represented as mean ± SEM. See also Figure 1.

**Figure S2.**
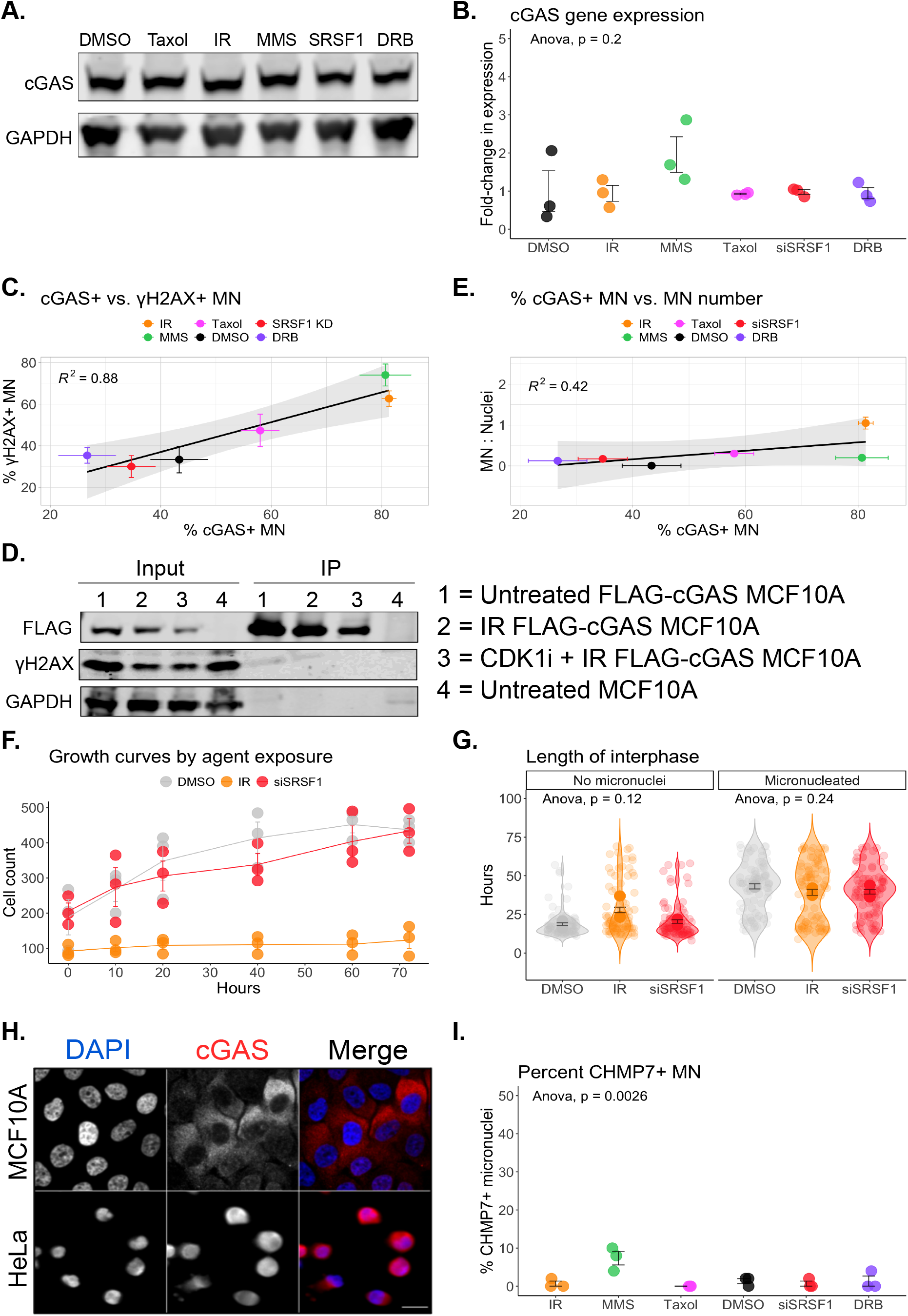
Variable recognition of MN across acute exposures is not an effect of MN number or changes to cGAS protein availability. (**A**) Western blot showing cGAS protein levels 72 hours post-acute exposures from MCF10A cells. Representative of 2 replicate experiments. (**B**) RT-qPCR for cGAS expression 72 hours post-acute exposures in MCF10A cells. Fold-change in expression is represented relative to average cGAS in the DMSO-treated samples. GAPDH was used as the housekeeping gene. Statistical comparison by two-way ANOVA. (**C**) Linear regression comparing the %γH2AX+ to the % cGAS+ MN observed by IF, 72 hours post-acute exposure. Each data point represents the mean of three biological replicates, where *n* per biological replicate = 50 MN. (**D**) FLAG-cGAS immunoprecipitated (IP) 72 hours following each of the four indicated treatment conditions, and probed with the indicated antibodies. CDK1 inhibitor was added one hour prior to acute exposure. Representative of 2 replicate experiments. (**E**) Linear regression comparing the number of MN in a microscopy field of view to the % cGAS+ MN observed by IF, 72 hours post-acute exposure. For % cGAS+ MN, each data point represents the mean of three biological replicates, where *n* per biological replicate = 50 MN. For MN: Nuclei, each data point represents the mean of the ratios observed in 3 biological replicates, *N* per biological replicate = 5-8 fields of view. (**F**) Growth curves for mCherry-H2B MCF10A cells in the 72 hours following each acute exposure. *N* per biological replicate = 80 – 500 cells. (**G**) The number of hours a cell spends in interphase. mCherry-H2B MCF10A cells observed for 72 hours post-acute exposures. *N* per biological replicate = 30 cells. (**H**) Representative IF images of untreated MCF10A and HeLa cells. Scale bar = 20 μm. (**I**)Percent CHMP7+ MN observed by IF, 72 hours post-acute exposure. *N* per biological replicate = 50 MN. Statistical comparison by two-way ANOVA. Data are represented as mean ± SEM. See also Figure 1.

**Figure S3.**
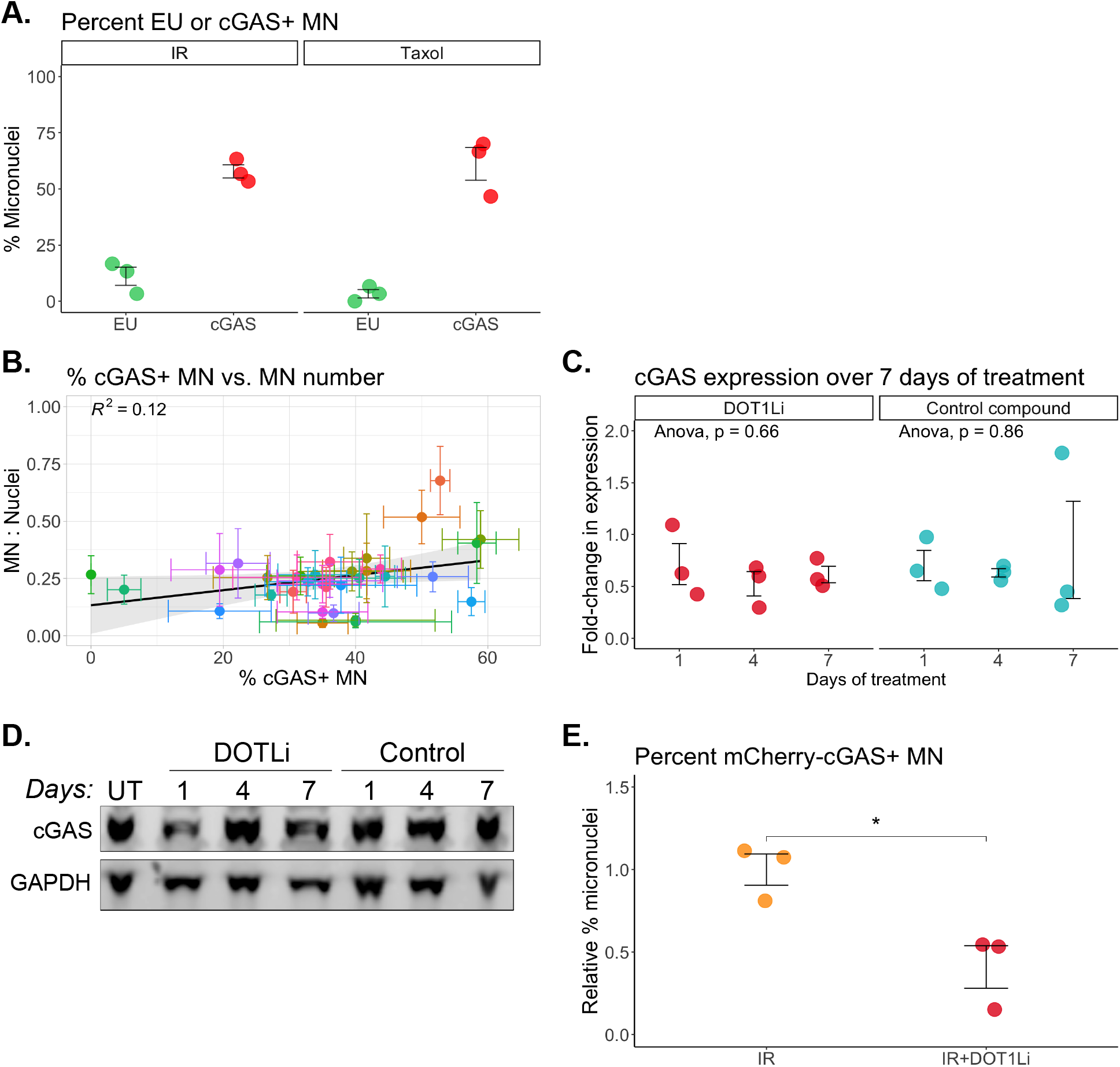
Validating the effects of chronic DOT1Li on % cGAS+ MN following IR. (**A**) Percent EU+ or cGAS+ MN observed in HeLa cells by IF, 72 hours post-acute exposure. *N* per biological replicate = 50 MN. (**B**) Linear regression comparing the number of MN in a microscopy field of view to the % cGAS+ MN observed by IF, 72 hours post-exposure to 10 Gy IR. Treatment with the library of pharmacological inhibitors began four days prior to IR exposure. Each colour represents a target listed in Fig. 3A and Table S1. For % cGAS+ MN, each data point represents the mean of three biological replicates, where *n* per biological replicate = 50 MN. For the ratio of MN to nuclei, each data point represents the mean of the ratios observed in 3 biological replicates, where *n* per biological replicate = 5 fields of view. (**C**) RT-qPCR for cGAS expression in MCF10A cells after one, four, and seven days of treatment with either DOT1Li (SGC0946) or its control compound (SGC0649). Fold-change in expression is represented relative to average cGAS in untreated samples. GAPDH was used as the housekeeping gene. Statistical comparison by two-way ANOVA. (**D**) Western blot for cGAS after one, four, and seven days of treatment with either DOT1Li or its control compound. Representative of 2 replicate experiments. (**E**) Percent mCherry-cGAS+ MN observed by IF, 72 hours post-exposure to 10 Gy IR in MCF10A cells. IR+DOT1Li cells were pre-treated with DOT1Li for four days prior to IR exposure. *N* per biological replicate = 50 MN. ns: *p* > 0.05, *: *p* <= 0.05, **: *p* <= 0.01, ***: *p* <= 0.001, ****: *p* <= 0.0001. Data are represented as mean ± SEM. See also Figure 2 and Figure 3.

**Figure S4.**
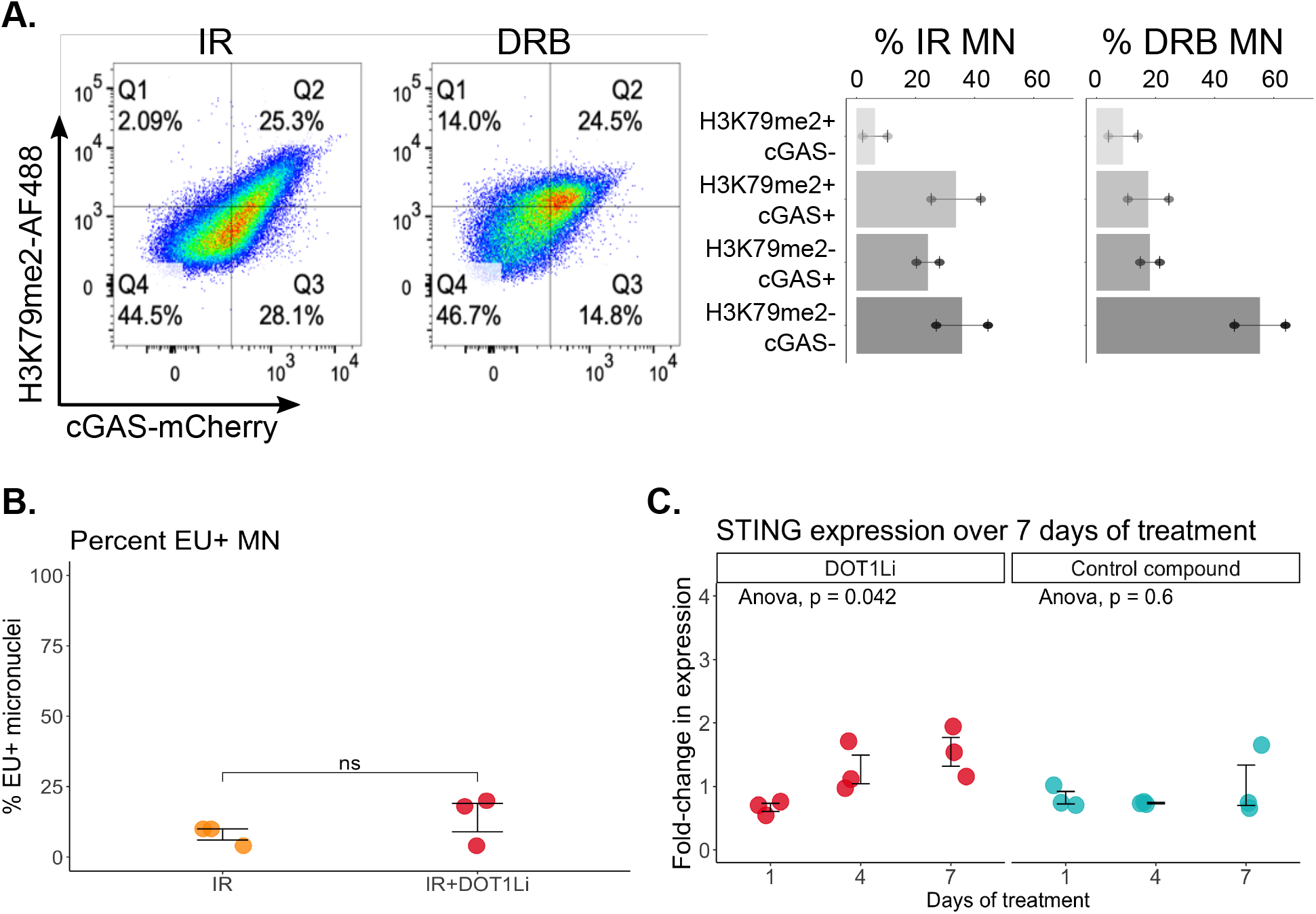
Active transcription in MN is not affected by changes to H3K79me2. (**A**) Flow cytometry plots depicting the relationship between H3K9me2+ and cGAS+ MN, in MN purified from IR- or DRB-treated HeLa cells. Gating scheme for MN is depicted in Figure S1A. Representative of 2 replicate experiments. (**B**) Percent EU+ MN observed by IF, 72 hours following IR. IR+DOT1Li cells were pre-treated with DOT1Li for four days prior to IR exposure. *N* per biological replicate = 50 MN. Statistical comparison by Student’s t-test. (**C**) RT-qPCR for STING expression after one, four, and seven days of treatment with either DOT1Li (SGC0946) or its control compound (SGC0649). Fold-change in expression is represented relative to average STING in untreated samples. GAPDH was used as the housekeeping gene. Statistical comparison by two-way ANOVA. ns: *p* > 0.05, *: *p* <= 0.05, **: *p* <= 0.01, ***: *p* <= 0.001, ****: *p* <= 0.0001. Data are represented as mean ± SEM. See also Figures 4 and 5.

## REFERENCES

Abe, T. and Barber, G.N. (2014) Cytosolic-DNA-Mediated, STING-Dependent Proinflammatory Gene Induction Necessitates Canonical NF-κB Activation through TBK1, Journal of Virology. Edited by B. Williams, 88(10), pp. 5328 LP – 5341. doi:10.1128/JVI.00037-141.

Bakhoum, M.F. et al. (2021) Loss of polycomb repressive complex 1 activity and chromosomal instability drive uveal melanoma progression., Nature communications, 12(1), p. 5402. doi:10.1038/s41467-021-25529-z2.

Bakhoum, S.F. et al. (2018) Chromosomal instability drives metastasis through a cytosolic DNA response, Nature, 553(7689), pp. 467–472. doi:10.1038/nature254323.

Barber, G.N. (2015) STING: infection, inflammation and cancer, Nature reviews. Immunology, 15(12), pp. 760–770. doi:10.1038/nri39214.

Chen, J., Harding, S.M., Natesan, R., Tian, L., Benci, J.L., Li, W., Minn, A.J., Asangani, I.A. and Greenberg, R.A. (2020) Cell Cycle Checkpoints Cooperate to Suppress DNA- and RNA-Associated Molecular Pattern Recognition and Anti-Tumor Immune Responses, Cell Reports, 32(9), p. 108080. doi:10.1016/j.celrep.2020.1080805.

Cohen S, Guenolé A, Marnef A, Clouaire T, Puget N, Rocher V, Arnould C, Aguirrebengoa M, Genais M, Vernekar D, Mourad R. Borde, L.G. (2020) BLM-dependent Break-Induced Replication handles DSBs in transcribed chromatin upon impaired RNA: DNA hybrids, bioRxiv 6.

Crasta, K. et al. (2012) DNA breaks and chromosome pulverization from errors in mitosis, Nature, 482(7383), pp. 53–58. doi:10.1038/nature108027.

Deng, L. et al. (2014) STING-Dependent Cytosolic DNA Sensing Promotes Radiation-Induced Type I Interferon-Dependent Antitumor Immunity in Immunogenic Tumors., Immunity, 41(5), pp. 843–852. doi:10.1016/j.immuni.2014.10.0198.

Flynn, P.J., Koch, P.D. and Mitchison, T.J. (2021) Chromatin bridges, not micronuclei, activate cGAS after drug-induced mitotic errors in human cells, Proceedings of the National Academy of Sciences. doi:10.1073/pnas.2103585118/-/DCSupplemental.Published9.

Gan, W., Guan, Z., Liu, J., Gui, T., Shen, K., Manley, J.L. and Li, X. (2011) R-loop-mediated genomic instability is caused by impairment of replication fork progression, Genes and Development, 25(19), pp. 2041–2056. doi:10.1101/gad.1701001110.

Gao, D., Wu, J., Wu, Y.-T., Du, F., Aroh, C., Yan, N., Sun, L. and Chen, Z.J. (2013) Cyclic GMP-AMP Synthase Is an Innate Immune Sensor of HIV and Other Retroviruses, Science, 341(6148), pp. 903 LP – 906. doi:10.1126/science.124093311.

Gentili, M. et al. (2019) The N-Terminal Domain of cGAS Determines Preferential Association with Centromeric DNA and Innate Immune Activation in the Nucleus, Cell Reports, 26(9), pp. 2377–2393.e13. doi:10.1016/j.celrep.2019.01.10512.

Harding, S.M., Benci, J.L., Irianto, J., Discher, D.E., Minn, A.J. and Greenberg, R.A. (2017) Mitotic progression following DNA damage enables pattern recognition within micronuclei, Nature, 548(7668), pp. 466–470. doi:10.1038/nature2347013.

Hatch, E.M., Fischer, A.H., Deerinck, T.J. and Hetzer, M.W. (2013) Catastrophic nuclear envelope collapse in cancer cell micronuclei, Cell, 154(1), pp. 47–60. doi:10.1016/j.physbeh.2017.03.04014.

Hatch, E.M.M., Fischer, A.H.H., Deerinck, T.J.J. and Hetzer, M.W.W. (2013) Catastrophic nuclear envelope collapse in cancer cell micronuclei, Cell, 154(1), pp. 47–60. doi:10.1016/j.physbeh.2017.03.04015.

Leibowitz, M.L., Papathanasiou, S., Doerfler, P.A., Blaine, L.J., Sun, L., Yao, Y., Zhang, C.Z., Weiss, M.J. and Pellman, D. (2021) Chromothripsis as an on-target consequence of CRISPR–Cas9 genome editing, Nature Genetics. doi:10.1038/s41588-021-00838-716.

Li, T., Huang, T., Du, M., Chen, X., Du, F., Ren, J. and Chen, Z.J. (2021) Phosphorylation and chromatin tethering prevent cGAS activation during mitosis, Science, 5386(February), pp. 1–1917.

Li, X.-D., Wu, J., Gao, D., Wang, H., Sun, L. and Chen, Z.J. (2013) Pivotal roles of cGAS-cGAMP signaling in antiviral defense and immune adjuvant effects., Science, 341(6152), pp. 1390–1394. doi:10.1126/science.124404018.

Liu, H. et al. (2018) Nuclear cGAS suppresses DNA repair and promotes tumorigenesis, Nature, 563(7729), pp. 131–136. doi:10.1038/s41586-018-0629-619.

Liu, S., Kwon, M., Mannino, M., Yang, N., Renda, F., Khodjakov, A. and Pellman, D. (2018) Nuclear envelope assembly defects link mitotic errors to chromothripsis, Nature, 561(7724), pp. 551–555. doi:10.1038/s41586-018-0534-z20.

Lutz, W.K., Tiedge, O., Lutz, R.W. and Stopper, H. (2005) Different types of combination effects for the induction of micronuclei in mouse lymphoma cells by binary mixtures of the genotoxic agents MMS, MNU, and genistein., Toxicological sciences: an official journal of the Society of Toxicology, 86(2), pp. 318–323. doi:10.1093/toxsci/kfi20021.

Ly, P., Teitz, L.S., Kim, D.H., Shoshani, O., Skaletsky, H., Fachinetti, D., Page, D.C. and Cleveland, D.W. (2017) Selective y centromere inactivation triggers chromosome shattering in micronuclei and repair by non-homologous end joining, Nature Cell Biology, 19(1), pp. 68–75. doi:10.1038/ncb345022.

MacDonald, K.M., Benguerfi, S. and Harding, S.M. (2020) Alerting the immune system to DNA damage: micronuclei as mediators, Essays in Biochemistry, 64(August), pp. 753–764. doi:10.1042/ebc2020001623.

Mackenzie, K.J. et al. (2017) cGAS surveillance of micronuclei links genome instability to innate immunity, Nature, 548(7668), pp. 461–465. doi:10.1038/nature2344924.

Maekawa, H. et al. (2019) Mitochondrial Damage Causes Inflammation via cGAS-STING Signaling in Acute Kidney Injury, Cell Reports, 29(5), pp. 1261–1273.e6. doi:10.1016/j.celrep.2019.09.05025.

Mammel, A.E., Huang, H.Z., Gunn, A.L., Choo, E. and Hatch, E.M. (2022) Chromosome length and gene density contribute to micronuclear membrane stability, Life Science Alliance, 5(2), p. e202101210. doi:10.26508/lsa.20210121026.

Michalski, S., de Oliveira Mann, C.C., Stafford, C., Witte, G., Bartho, J., Lammens, K., Hornung, V. and Hopfner, K.P. (2020) Structural basis for sequestration and autoinhibition of cGAS by chromatin, Nature, (May). doi:10.1038/s41586-020-2748-027.

Mohr, L., Toufektchan, E., von Morgen, P., Chu, K., Kapoor, A. and Maciejowski, J. (2021) ER-directed TREX1 limits cGAS activation at micronuclei, Molecular Cell, 0(0), pp. 1–15. doi:10.1016/j.molcel.2020.12.03728.

Nguyen, A.T., Taranova, O., He, J. and Zhang, Y. (2011) DOT1L, the H3K79 methyltransferase, is required for MLL-AF9-mediated leukemogenesis, Blood. 2011/04/26, 117(25), pp. 6912–6922. doi:10.1182/blood-2011-02-33435929.

Oksenych, V. et al. (2013) Histone Methyltransferase DOT1L Drives Recovery of Gene Expression after a Genotoxic Attack, PLoS Genetics, 9(7). doi:10.1371/journal.pgen.100361130.

Pathare, G.R. et al. (2020) Structural mechanism of cGAS inhibition by the nucleosome, Nature, (February). doi:10.1038/s41586-020-2750-631.

Qing Chen, Adrienne Boire, Xin Jin, Manuel Valiente, E.E.E. and A.L.-S. (2016) Carcinoma-astrocyte gap junctions promote brain metastasis by cGAMP transfer, Nature, 53332.

Scheer, S. et al. (2019) A chemical biology toolbox to study protein methyltransferases and epigenetic signaling, Nature Communications, 10(1), p. 19. doi:10.1038/s41467-018-07905-433.

Sepaniac, L.A., Martin, W., Dionne, L.A., Stearns, T.M., Reinholdt, L.G. and Stumpff, J. (2021) Micronuclei in Kif18a mutant mice form stable micronuclear envelopes and do not promote tumorigenesis, Journal of Cell Biology, 220(11) 34.

Shanbhag, N.M. and Greenberg, R.A. (2013) The Dynamics of DNA Damage Repair and Transcription, Methods in Molecular Biology, 103(2), pp. 1–9. doi:10.1159/00009062035.

Shanbhag, N.M., Rafalska-Metcalf, I.U., Balane-Bolivar, C., Janicki, S.M. and Greenberg, R.A. (2010) ATM-dependent chromatin changes silence transcription in cis to DNA doublestrand breaks., Cell, 141(6), pp. 970–981. doi:10.1016/j.cell.2010.04.03836.

Shimizu, N., Kanda, T. and Wahl, G.M. (1996) Selective capture of acentric fragments by micronuclei provides a rapid method for purifying extrachromosomally amplified DNA., Nature Genetics, 12(1), pp. 65–71. doi:10.1038/ng0196-6537.

Sidoli, S., Bhanu, N. V., Karch, K.R., Wang, X. and Garcia, B.A. (2016) Complete workflow for analysis of histone post-translational modifications using bottom-up mass spectrometry: From histone extraction to data analysis, Journal of Visualized Experiments, 2016(111), pp. 1–11. doi:10.3791/5411238.

Sommer, S., Buraczewska, I. and Kruszewski, M. (2020) Micronucleus Assay: The State of Art, and Future Directions, International Journal of Molecular Sciences, pp. 7–9. doi:10.3390/ijms2104153439.

Tang, J., Cho, N.W., Cui, G., Manion, E.M., Shanbhag, N.M., Botuyan, M.V., Mer, G. and Greenberg, R.A. (2013) Acetylation limits 53BP1 association with damaged chromatin to promote homologous recombination., Nature structural & molecular biology, 20(3), pp. 317–325. doi:10.1038/nsmb.249940.

Terradas, M., Martín, M. and Genescà, A. (2016) Impaired nuclear functions in micronuclei results in genome instability and chromothripsis, Archives of Toxicology, 90(11), pp. 2657–2667. doi:10.1007/s00204-016-1818-441.

Terradas, M., Martín, M., Tusell, L. and Genescà, A. (2010) Genetic activities in micronuclei: Is the DNA entrapped in micronuclei lost for the cell?, Mutation Research - Reviews in Mutation Research, 705(1), pp. 60–67. doi:10.1016/j.mrrev.2010.03.00442.

Tigano, M., Vargas, D.C., Tremblay-Belzile, S., Fu, Y. and Sfeir, A. (2021) Nuclear sensing of breaks in mitochondrial DNA enhances immune surveillance, Nature, 591(7850), pp. 477–481. doi:10.1038/s41586-021-03269-w43.

Uggenti, C. et al. (2020) cGAS-mediated induction of type I interferon due to inborn errors of histone pre-mRNA processing, Nature Genetics. doi:10.1038/s41588-020-00737-344.

Umbreit, N.T. et al. (2020) Mechanisms Generating Cancer Genome Complexity From A Single Cell Division Error, Science, 368(6488), p. 835058. doi:10.1101/83505845.

Utani, K.I., Kohno, Y., Okamoto, A. and Shimizu, N. (2010) Emergence of micronuclei and their effects on the fate of cells under replication stress, PLoS ONE, 5(4). doi:10.1371/journal.pone.001008946.

Vanpouille-Box, C. et al. (2017) DNA exonuclease Trex1 regulates radiotherapy-induced tumour immunogenicity., Nature communications, 8, p. 15618. doi:10.1038/ncomms1561847.

Vietri, M., Schultz, Sebastian W, et al. (2020) Unrestrained ESCRT-III drives micronuclear catastrophe and chromosome fragmentation, Nature Cell Biology. doi:10.1038/s41556-020-0537-548.

Vietri, M., Schultz, Sebastian W., et al. (2020) Unrestrained ESCRT-III drives micronuclear catastrophe and chromosome fragmentation, Nature Cell Biology, 22(7), pp. 856–867. doi:10.1038/s41556-020-0537-549.

Volkova, N. V. et al. (2020) Mutational signatures are jointly shaped by DNA damage and repair, Nature Communications, 11(1). doi:10.1038/s41467-020-15912-750.

Wan, L. et al. (2021) Translation stress and collided ribosomes are co-activators of cGAS, Molecular Cell, pp. 1–15. doi:10.1016/j.molcel.2021.05.01851.

West, A.P. et al. (2015) Mitochondrial DNA stress primes the antiviral innate immune response, Nature, 520(7548), pp. 553–557. doi:10.1038/nature1415652.

Wu, Q. et al. (2019) A chemical toolbox for the study of bromodomains and epigenetic signaling., Nature communications, 10(1), p. 1915. doi:10.1038/s41467-019-09672-253.

Xu, B. et al. (2011) Replication stress induces micronuclei comprising of aggregated DNA double-strand breaks, PLoS ONE, 6(4). doi:10.1371/journal.pone.001861854.

Yu, W. et al. (2012) Catalytic site remodelling of the DOT1L methyltransferase by selective inhibitors, Nature Communications, 3, pp. 1–12. doi:10.1038/ncomms230455.

Zhang, C.Z., Spektor, A., Cornils, H., Francis, J.M., Jackson, E.K., Liu, S., Meyerson, M. and Pellman, D. (2015) Chromothripsis from DNA damage in micronuclei, Nature, 522(7555), pp. 179–184. doi:10.1038/nature1449356.

Zhao, B. et al. (no date) Topoisomerase 1 cleavage complex enables pattern recognition and inflammation during senescence, Nature Communications, (2020). doi:10.1038/s41467-020-14652-y57.

Zhao, B., Xu, P., Rowlett, C.M., Jing, T., Shinde, O., Lei, Y., West, A.P., Liu, W.R. and Li, P. (2020) The Molecular Basis of Tight Nuclear Tethering and Inactivation of cGAS, Nature, (May). doi:10.1038/s41586-020-2749-z58.

Zhou, W., Mohr, L., Maciejowski, J., Kranzusch, P.J., Zhou, W., Mohr, L., Maciejowski, J. and Kranzusch, P.J. (2021) cGAS phase separation inhibits TREX1-mediated DNA degradation and enhances cytosolic DNA sensing, Molecular Cell, 81(4), pp. 739–755.e7. doi:10.1016/j.molcel.2021.01.02459.

